# VO659, a CAG repeat-targeting antisense oligonucleotide, lowers multiple neurotoxic mutant huntingtin proteins in Huntington’s disease

**DOI:** 10.64898/2026.10.16.758452

**Authors:** Bas Röttgering, Rudie Weij, Suzanne Bijl, Ruurd Verheul, Janwillem Testerink, Lodewijk Toonen, Robert M. Bragg, Jeffrey B. Carroll, Chantal Beekman, Nicole Datson

**Author notes:** Correspondence should be addressed to N.D., Address: J.H. Oortweg 21, 2333 CH Leiden, The Netherlands.

## Abstract

Huntington’s disease (HD) is caused by a CAG repeat expansion in the huntingtin (*HTT*) gene, encoding an elongated polyglutamine stretch in the corresponding mutant HTT (mHTT) protein. HD pathogenesis is driven by multiple neurotoxic mHTT protein species, including HTT1a and repeat-associated non-AUG translation (RAN) proteins, which increase in expression as mutant CAG repeats lengthen due to somatic expansion. VO659 is an investigational CAG repeat-targeting antisense oligonucleotide (ASO) that lowers mHTT protein levels by blocking translation. Here, we show that VO659 allele-preferentially lowers levels of multiple neurotoxic mHTT protein species in a CAG repeat length-dependent manner, relatively sparing wild-type HTT, while becoming more efficacious as somatic expansion occurs. VO659 reduced *HTT1a* mRNA levels in HD fibroblasts with an ultra-long CAG repeat, blocked the translation of HTT1a and polyalanine-containing RAN proteins derived from *HTT* exon 1 overexpression constructs, and displayed a long tissue half-life in mouse brain, supporting an infrequent dosing regimen. A first-in-human, open-label phase 1/2a clinical trial (NCT05822908) is currently ongoing in HD patients. VO659’s unique CAG repeat-dependent and allele-preferential mechanism of action holds promise for its therapeutic effect in HD patients by targeting multiple core neurotoxic proteins derived from (somatically) expanded *mHTT* CAG repeats, while sparing wild-type HTT expression.

## Introduction

Huntington’s Disease (HD) is a rare, dominantly inherited, progressive and fatal neurodegenerative disorder caused by a CAG trinucleotide repeat expansion in exon 1 of the huntingtin (*HTT*) gene.^1^ In non-affected individuals the CAG repeat in *HTT* is between 6–35 triplets, while in HD patients the repeat is expanded to ≥40 triplets. Intermediate CAG repeats of 36–39 result in reduced disease penetrance and repeats of ≥56 triplets often result in juvenile-onset HD.^2^ Due to neurodegeneration, patients with HD progressively develop motor symptoms, including chorea, dysphagia, and loss of ambulation, as well as cognitive and psychiatric symptoms such as executive dysfunction, apathy, and depression.^3–4^ Treatment of HD is currently limited to symptom management, as there is no approved disease-modifying treatment yet available.

There are multiple pathogenic mechanisms described through which the CAG repeat expansion in *HTT* leads to neurodegeneration. Historically, the primary pathogenic mechanism was thought to be the translation of the mutant CAG repeat into an extended polyglutamine (polyQ) tract that confers a toxic gain-of-function to the mutant HTT (mHTT) protein. Expression of the mHTT protein leads to disruption of several processes, including transcriptomic dysregulation and impaired mitochondrial metabolism.^5–7^ Additionally, the extended polyQ tract makes the mHTT protein aggregation-prone,^8^ leading to detectable mHTT aggregates in various brain regions in post-mortem brains of HD patients, including in the striatum and cerebral cortex.^9^ Furthermore, proteolytic cleavage of the full-length mHTT protein generates toxic and aggregation-prone shorter N-terminal protein fragments.^10–13^ Besides toxic gain-of-function of mHTT, partial loss-of-function of wild-type HTT (wtHTT) may also contribute to HD pathogenesis.^14^

Recently, somatic instability, leading to expansion of the germline inherited mutant CAG repeat, has been shown to be an important proximal driver of neurotoxicity.^15–16^ It is thought that, initially, expansion up to 150 CAGs occurs slowly over the course of decades, after which the rate of expansion increases, leading to severe transcriptional dysregulation, neuronal dysfunction and ultimately neuronal death.^17–18^ Although somatic expansion occurs throughout the HD brain,^19^ the extent of somatic expansion may be cell-type dependent. Using single cell RNA profiling of post-mortem HD brains, it was shown that striatal projection neurons exhibit the most pronounced somatic expansion, while in the cortex glutamatergic neurons show the highest rate of expansion in both cases resulting in ultra-long CAG repeats exceeding 150– 500 CAGs.^17–18^

The ultra-long CAG repeats arising from somatic expansion result in the formation of additional toxic HTT protein species, including HTT1a (Fig. 1). HTT1a is extremely aggregation-prone and possibly the most neurotoxic mHTT protein,^20–22^ suggesting that HTT1a is one of the most important drivers of HD pathology and therefore an important therapeutic target. HTT1a is encoded by the mis-spliced *HTT1a* transcript comprising only *HTT* exon 1 and part of intron 1, terminating at a cryptic polyadenylation site ∼7 kb into intron 1.^23–26^ Formation of *HTT1a* is CAG repeat length-dependent,^26^ but the precise molecular mechanisms leading to *HTT1a* production are not completely understood, although aberrant binding of splicing factors to the CAG repeat, aberrant methylation of adenine bases (m^6^A) in *HTT* pre-mRNA, aberrant export of *HTT* exon1-intron1 mRNA from the nucleus, and the proline-rich domain in exon 1 of the *HTT* transcript may play a role.^24–25,27–30^

**Figure 1.**
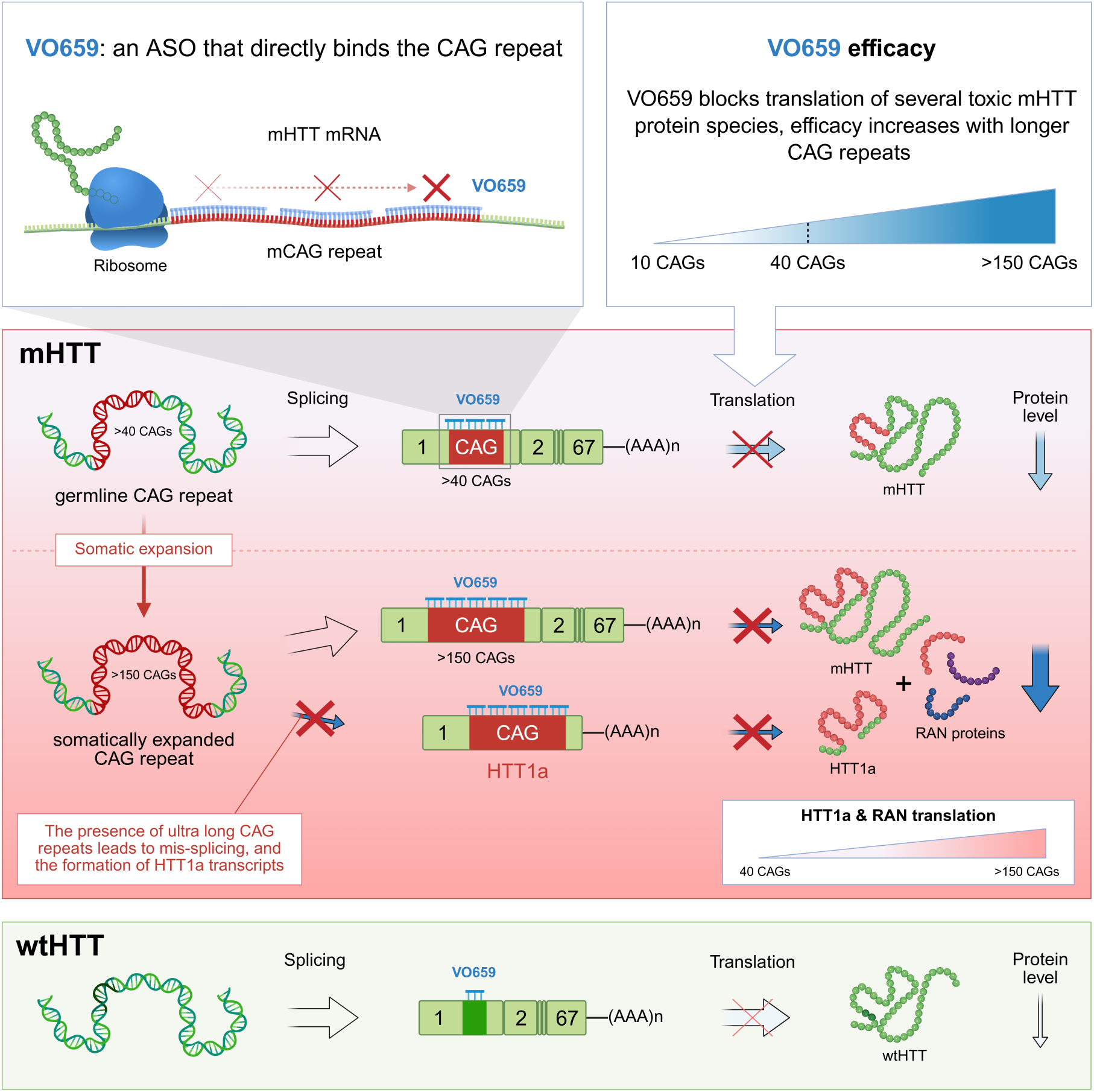
VO659 targets multiple mHTT-derived toxic protein species. VO659, a fully modified 2’-*O*-methyl (2’-*O*-Me) phosphorothioate (PS) antisense oligonucleotide (ASO), binds to the CAG repeat in exon 1 of *HTT* RNA. More VO659 molecules can bind as the CAG repeat lengthens, leading to more effective hindrance of translation on longer repeats. This underlies the allele-preferential lowering of HTT protein levels. When the CAG repeat in the *HTT* gene undergoes somatic expansion, VO659 becomes even more efficacious, also reducing levels of additional *mHTT*-derived protein species, including toxic HTT1a and repeat-associated non-AUG (RAN) translation-derived proteins. VO659 can lower HTT1a levels in 2 ways: it can inhibit mutant CAG repeat-related mis-splicing that results in toxic *HTT1a* transcripts when binding the CAG repeat in *HTT* pre-mRNA. In addition, VO659 can bind the remaining *HTT1a* mRNA and hinder its translation. Similarly, VO659 binds to the expanded CAG repeat and prevents RAN translation of toxic RAN proteins.

Furthermore, expanded CAG repeats in *mHTT* mRNA have been shown to form secondary structures and hairpins that can induce repeat-associated non-AUG (RAN) translation, a non-canonical form of translation that initiates within the expanded repeat.^31–33^ RAN translation leads to the production of polyglutamine (polyQ) and frameshifted polyserine (polyS) and polyalanine (polyA) proteins from the three reading frames of *mHTT* mRNA, as well as polyleucine (polyL), polycysteine (polyC), and polyalanine (polyA) proteins from the mutant *HTT antisense* (*HTT-AS*) transcript.^34^ Expression of these short repeat proteins has been shown to be toxic *in vitro* and analysis of post-mortem brains of HD patients has identified polyS, polyA, polyL and polyC expression and aggregation in affected brain regions that display neuronal loss, microglial activation and apoptosis, including the striatum, frontal cortex and cerebellum.^34–35^ Expression of RAN proteins has been shown to increase with CAG repeat length,^34^ suggesting that their production increases as somatic expansion occurs. Overall, this indicates that expression and aggregation of the full-length mHTT protein is likely not the sole driver of HD pathogenesis. As mutant CAG repeats become progressively longer through somatic expansion, additional toxic mHTT protein species are formed, including HTT1a and RAN proteins, that drive disease onset, neurodegeneration, and clinical progression in HD.

Here we show that VO659, an investigational clinical stage steric blocking ASO that directly targets the disease-causing CAG repeat mutation in HD, possesses a unique mechanism of action (MoA) relevant to multiple molecular drivers of disease pathophysiology. We show that VO659 acts in a CAG repeat length-dependent and thus allele-preferential manner, relatively sparing important wtHTT levels while reducing levels of mHTT protein with increased efficacy as CAG repeats expand. We demonstrate that VO659 efficiently lowers levels of different toxic mHTT protein species, including the HTT1a isoform and polyA-containing RAN proteins that increase in expression upon somatic CAG repeat expansion, thus targeting multiple CAG repeat-related drivers of neurotoxicity in HD (Fig. 1). Furthermore, we demonstrate that VO659 has a favorable pharmacokinetic (PK) profile upon intracerebroventricular (ICV) dosing in mice, with good uptake across the central nervous system (CNS) and a long tissue half-life, making it suitable for infrequent dosing. VO659 is the only clinical stage (NCT05822908) HTT-lowering therapy to date with the potential to allele-preferentially lower full-length mHTT protein levels while simultaneously targeting all currently described *mHTT*-derived drivers of neurotoxicity in HD and therefore holds promise for the treatment of HD.

## Results

### Targeting the CAG repeat expansion with VO659 lowers HTT protein levels in a CAG repeat length-dependent and allele-preferential manner

The CAG repeat-dependent efficacy of VO659 was tested in a panel of HD patient-derived fibroblasts with various mutant repeat lengths ranging from 43 to 175 CAGs and in a healthy control fibroblast line. The fibroblasts were transfected with 25–100 nM of VO659 or a non-allele-selective *HTT* exon 36-targeting RNase-H recruiting gapmer ASO (HTT-gap36) with the same sequence and chemistry as HTT_rx_, also known as tominersen, as a comparator.^36–37^ To facilitate size separation of wtHTT and mHTT proteins, we applied a previously described caspase 3 digestion-based Simple Western (SW) capillary immunoassay to generate N-terminal wtHTT and mHTT protein fragments containing the polyQ region that could be size-separated and quantified (Fig. 2A–C).^38^ VO659 inhibited translation of *HTT* mRNA, leading to decreased HTT protein levels with a clear CAG repeat length-dependent effect (Fig. 2D). While wtHTT protein levels showed almost no reduction by VO659 up to 50 nM and reached a maximum reduction of 29% at 100 nM, mHTT protein levels were reduced more profoundly by VO659 (Fig. 2A–D). In HD patient-derived fibroblasts with adult-onset repeat lengths of 43– 45 CAGs, VO659 reduced mHTT protein levels dose-dependently, with an average reduction of 40% at a concentration of 100 nM (Fig. 2A, D). In a juvenile HD patient fibroblast line with 68 CAG repeats, the average reduction of mHTT protein was 50% at the highest concentration (Fig. 2B, D) and further increased to approximately 75% in a cell line with an ultra-long repeat of 175 CAGs (Fig. 2C, D).

**Figure 2.**
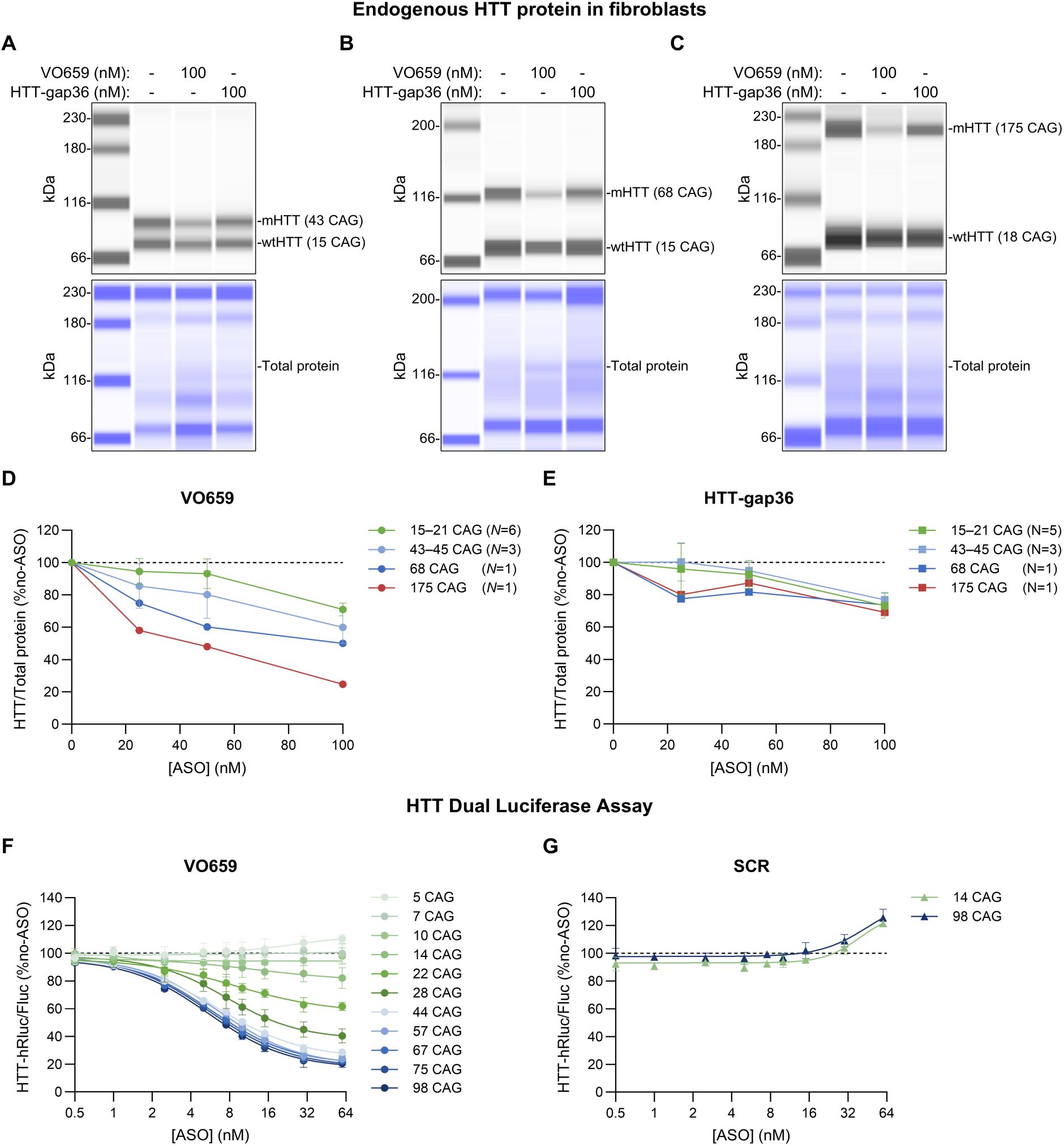
VO659 reduces HTT protein levels in a CAG repeat length-dependent manner **A–C** Representative Simple Western (SW) blot images of HD fibroblast cell lines with 43:15 **(A)**, 68:15 **(B)**, or 175:18 **(C)** CAG repeats transfected with 100 nM VO659 or HTT-gap36. To facilitate size separation of wtHTT and mHTT, protein lysates were treated with caspase-3 prior to SW analysis using an N-terminal anti-HTT antibody (Abcam, ab109115, 1:100).^38^ Total protein was used for normalization. **D– E** Quantification of HTT protein levels measured by SW analysis in VO659-**(D)** and HTT-gap36-**(E)** treated samples, normalized to total protein. *N* indicates the number of measured cell lines per range of CAG repeat length. Data are expressed as mean ± standard deviation (SD) and as a percentage of no-ASO controls (set at 100% per CAG repeat). **F–G** Dual-Luciferase reporter (DLR) assay with VO659 **(F)** or scrambled (SCR) **(G)**. HeLa cells were transfected with ASOs followed by plasmids with 5–98 CAG repeats. HTT-Renilla fusion reporter (HTT-hRluc) activity was normalized to Firefly (Fluc) internal control activity and expressed as % of no-ASO controls per plasmid. Data represent mean ± SD of 3 independent transfections. Non-linear regression lines (four parameter – variable slope) were fitted to the data using GraphPad Prism.

In contrast, when the HD fibroblast lines were transfected with increasing concentrations of the non-allele-preferential HTT-gap36 ASO, no allele-preferential HTT protein knockdown was observed (Fig. 2E). HTT-gap36 reduced wtHTT protein levels by 27% across all cell lines at 100 nM, while mHTT protein at 100 nM was reduced by 23%, 26% and 31% in HD fibroblast lines carrying 43–45, 68 or 175 CAG repeats, respectively. When 25–100 nM of a scrambled version of VO659 (SCR) was transfected into the HD fibroblast line with 175 and 18 CAG repeats in m*HTT* and *wtHTT* respectively, no effect on mHTT or wtHTT protein levels was observed (Supplemental Fig. S1).

The CAG repeat length-dependent translational inhibition of VO659 was additionally characterized using a dual luciferase reporter (DLR) assay. DLR constructs were generated in which *HTT* exons 1–6 with increasing CAG repeat lengths (5–98 repeats) were fused to the 5′ end of *Renilla* luciferase (*hRluc*) gene. The vector also contained a Firefly luciferase (*Fluc*) internal control signal driven by a separate promoter. These constructs were transfected into HeLa cells together with 0.5–60 nM of VO659 (Fig. 2F). Similar to the fibroblast data, VO659 showed a clear CAG repeat length-dependent inhibition of translation, leading to reduced activity of the HTT-hRluc reporter. With constructs containing up to 10 CAG repeats, no clear dose-dependent reduction in HTT-hRluc reporter activity was observed. VO659 induced a limited dose-dependent reduction in HTT-hRluc reporter activity of the construct with 14 CAG repeats, with a maximum reduction of 18% at 60 nM. With constructs containing 22 and 28 CAGs, the reduction at 60 nM VO659 increased to 38% and 60%, respectively. In the mutant CAG repeat range, VO659 induced larger dose-dependent reductions in HTT-hRluc reporter signal. In constructs with 44 CAG repeats, the maximum observed reduction was 72% at 60 nM VO659. A drawback of the DLR assay under the used experimental conditions was that it reached a functional plateau around an ∼80% reduction in reporter activity, offering only limited resolution to detect increases in efficacy of VO659 between longer (e.g. 57–98) CAG repeats.

To verify that the CAG repeat-dependent reductions were caused by direct CAG repeat targeting and not by any hybridization independent ASO effects, 0.5–60 nM of a scrambled (SCR) version of VO659 was transfected together with the DLR constructs. As expected, no reduction in HTT-hRluc reporter activity was induced by the SCR ASO regardless of CAG repeat length, although a moderate increase in luciferase signal was observed at 60 nM SCR (Fig. 2G).

Together, these data derived from two independent assays demonstrate that VO659 reduces HTT protein levels in a CAG repeat length-dependent manner, and that the efficacy of VO659 further increases when binding to juvenile HD onset-range and ultra-long (175 CAG) repeats in fibroblasts.

### VO659 does not directly impact somatic expansion, but acts on its downstream drivers of neurotoxicity

Having shown that VO659 reduces HTT protein levels in a CAG repeat-dependent manner, and thus becomes more efficacious as somatic expansion occurs, we next examined whether VO659 may also directly reduce the rate of somatic expansion. Chronic treatment with a gapmer ASO acting in the nucleus has been shown to repress *Htt* transcription in HD mouse models, leading to significantly reduced somatic expansion *in vivo*. In contrast, siRNAs that act in the cytoplasm did not affect somatic expansion despite lowering Htt protein levels.^39^ VO659 has a mechanism of action distinct from these examples as it binds the CAG repeat in *mHTT* and blocks its translation in the cytoplasm, but can also enter the nucleus and bind pre-mRNA. To examine whether VO659 could prevent somatic expansion, Q111 mice^40^ received a total of 4 ICV doses of VO659, once every 2 weeks, and were sacrificed 6 weeks after the last dose, after which somatic expansion was assessed in the striatum. No effect of VO659 treatment was observed on somatic expansion, despite a significant reduction of mHTT protein of 20% compared to vehicle (VEH) in the cortex (Supplemental Fig. S2). As expected, no reduction of wtHTT protein was observed given the short and interrupted CAG repeat sequence in the wild-type allele of the Q111 mouse ((CAG)_2_CAA(CAG)_4_).

While VO659 did not directly affect somatic expansion, its translational inhibition of full-length *mHTT* transcripts with ultra-long CAG repeats resulting from somatic expansion is expected to be highly effective, as we have observed in HD fibroblasts with 175 CAGs (Figure 2D).

### VO659 lowers HTT1a protein levels through translation inhibition

One of the expected downstream consequences of somatic expansion is the production of the *HTT* exon 1-derived *HTT1a* transcript that arises from alternative processing or mis-splicing of *mHTT* pre-mRNA containing very long CAG repeats, in which a cryptic polyadenylation site in intron 1 is utilized.^23–25^ Since VO659 targets the CAG repeat in exon 1 of *mHTT,* we investigated whether VO659 could inhibit translation of the *HTT1a* transcript to protein. Considering *HTT1a* mRNA is only expressed at very low levels in HD fibroblasts and protein would therefore be difficult to detect,^26^ we transfected HeLa cells with plasmids containing an artificial *HTT1a* overexpression construct with 100 or 156 CAG repeats, and a GFP internal control driven by a separate promoter to normalize for transfection efficacy. HTT1a was detected with immunostaining using a HTT1a-specific antibody (#CHDI-90004291 – 11G2).^41^ HTT1a could only clearly be detected in cells successfully transfected with plasmid, as indicated by GFP positivity, confirming the specificity of the antibody (Fig. 3A). Quantification of HTT1a intensity showed that VO659 dose-dependently decreased HTT1a levels (Fig. 3B). In cells transfected with the 100 or 156 CAG constructs, 50 nM VO659 decreased HTT1a intensity by 68% and 66%, respectively. Interestingly, high-intensity puncta could be observed (Fig. 3A, white arrows) in some GFP-positive (GFP+) cells, which increased in frequency with CAG repeat length (Fig. 3C). These puncta most likely represent HTT1a aggregates, since it is known that HTT1a is very aggregation prone.^21–22,42–43^ VO659 dose-dependently reduced the percentage of GFP+ cells that contained HTT1a aggregates, resulting in a lowering at 50 nM from 9.3% to 2.1% in cells expressing the 100 CAG construct, and from 13.2% to 3.6% in cells expressing the 156 CAG HTT1a overexpression construct (Fig. 3C). In contrast, transfection with 50 nM of non-targeting HTT-gap36 or SCR had no effect on HTT1a intensity or HTT1a aggregates.

**Figure 3.**
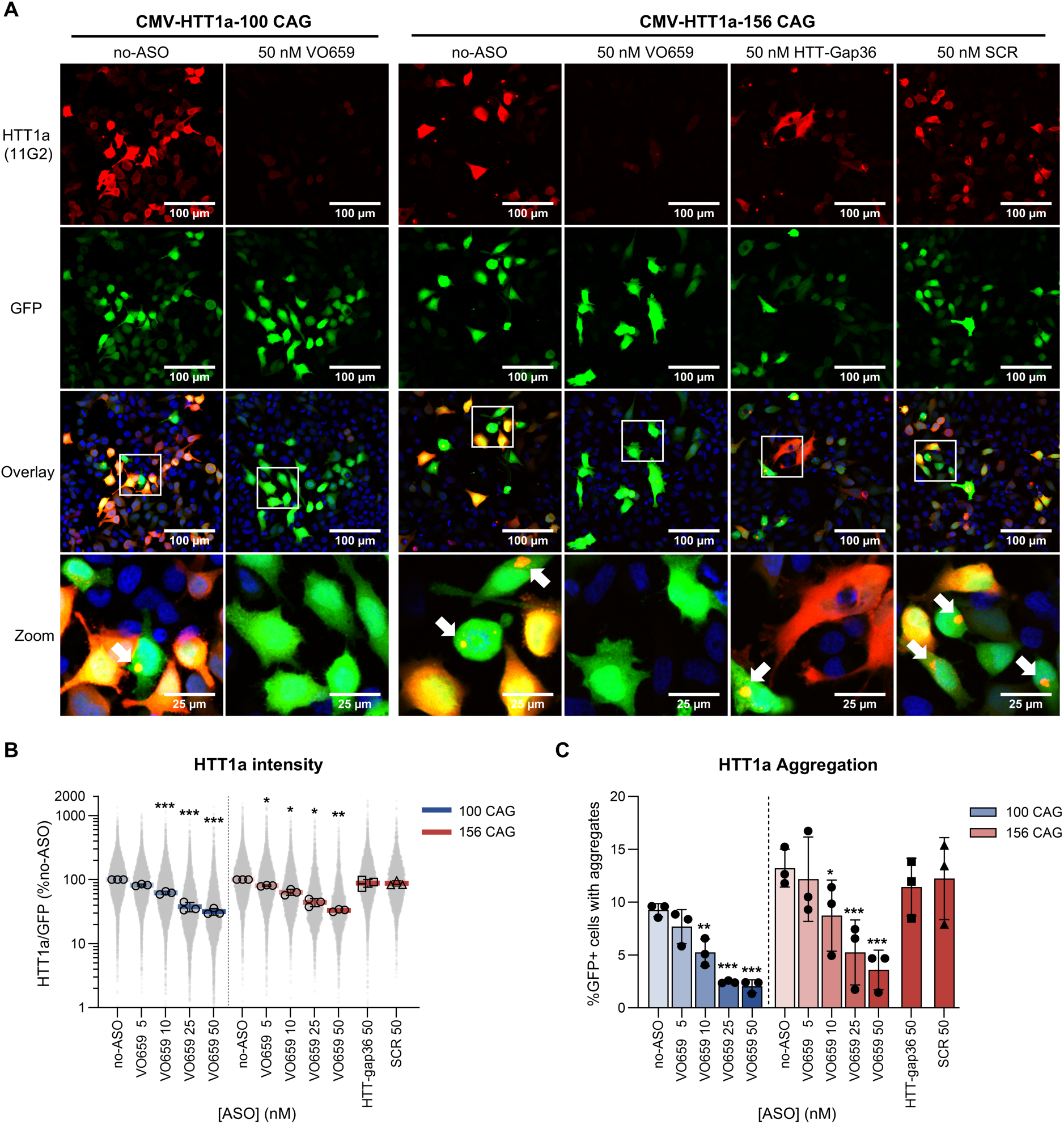
VO659 sterically hinders translation of HTT1a **A** Representative 20x confocal images of HeLa cells transfected with HTT1a overexpression constructs containing 100 or 156 CAG repeats (co-expressing GFP from a separate promoter) and 50 nM VO659, HTT-gap36, or SCR. Green: GFP; Red: HTT1a (CHDI foundation, #CHDI-90004291 – 11G2, 1:500)^41^; Blue: DAPI nuclear counterstain (1:1000). White boxes indicate the digitally zoomed areas shown in the bottom row; white arrows indicate HTT1a aggregates. **B** HTT1a staining intensity in HeLa cells quantified with high content imaging, corrected for GFP intensity. Colored horizontal lines represent the mean ± SD of three independent experiments (The no-ASO control was set at 100% per plasmid). Open symbols depict the average per independent experiment. The grey data points represent the normalized HTT1a intensity per cell, pooled from all experiments. A total of 107,775 GFP+ cells were analyzed. The y-axis minimum was set at 1% for visualization, resulting in the clipping of 41 cells from the graph (these cells were still included in calculations). **C** The percentage of GFP+ HeLa cells (e.g. cells successfully transfected with the HTT1a overexpression constructs) containing at least one HTT1a aggregate, quantified with high content imaging, from the same three experiments as in B. The average ± SD is plotted. Groups were compared to the no-ASO condition using a one-way repeated-measures ANOVA with a post-hoc Dunnett’s test in GraphPad Prism. For B, statistics were performed on non-normalized log-transformed data. * = p<0.05, ** = p<0.005, *** = p<0.0005

### VO659 lowers *HTT1a* mRNA levels through preventing mis-splicing

Having demonstrated that VO659 can inhibit translation and thus lower levels of HTT1a protein, we next investigated whether VO659 could also prevent the mis-splicing leading to formation of *HTT1a* mRNA. It has been suggested that proteins aberrantly binding *mHTT* pre-mRNA due to the expanded CAG repeat may lead to mis-splicing and *HTT1a* expression.^24–25,28–29^ We hypothesized that this process could be sterically hindered by VO659 binding the CAG repeat, leading to lower *HTT1a* mRNA levels.

To examine this, *HTT1a* mRNA levels were quantified using a reverse transcriptase digital PCR (RT-dPCR) assay as described by Hoschek et al.^26^ (Fig. 4A) in HD patient fibroblasts. As previously reported, *HTT1a* expression was only detectable at low levels in HD patient fibroblasts with ultra-long (175) CAG repeats (Fig. 4B).^26^ *HTT1a* mRNA could not be reliably detected in HD fibroblast lines carrying 68 or 43 CAG repeats in *mHTT*. Similarly, *HTT1a* levels were undetectable in a fibroblast line from another polyQ expansion disorder, spinocerebellar ataxia type 1 (SCA1), which carries non-pathogenic *HTT* alleles with 18 CAG repeats on both *wtHTT* alleles (Fig. 4B).

**Figure 4.**
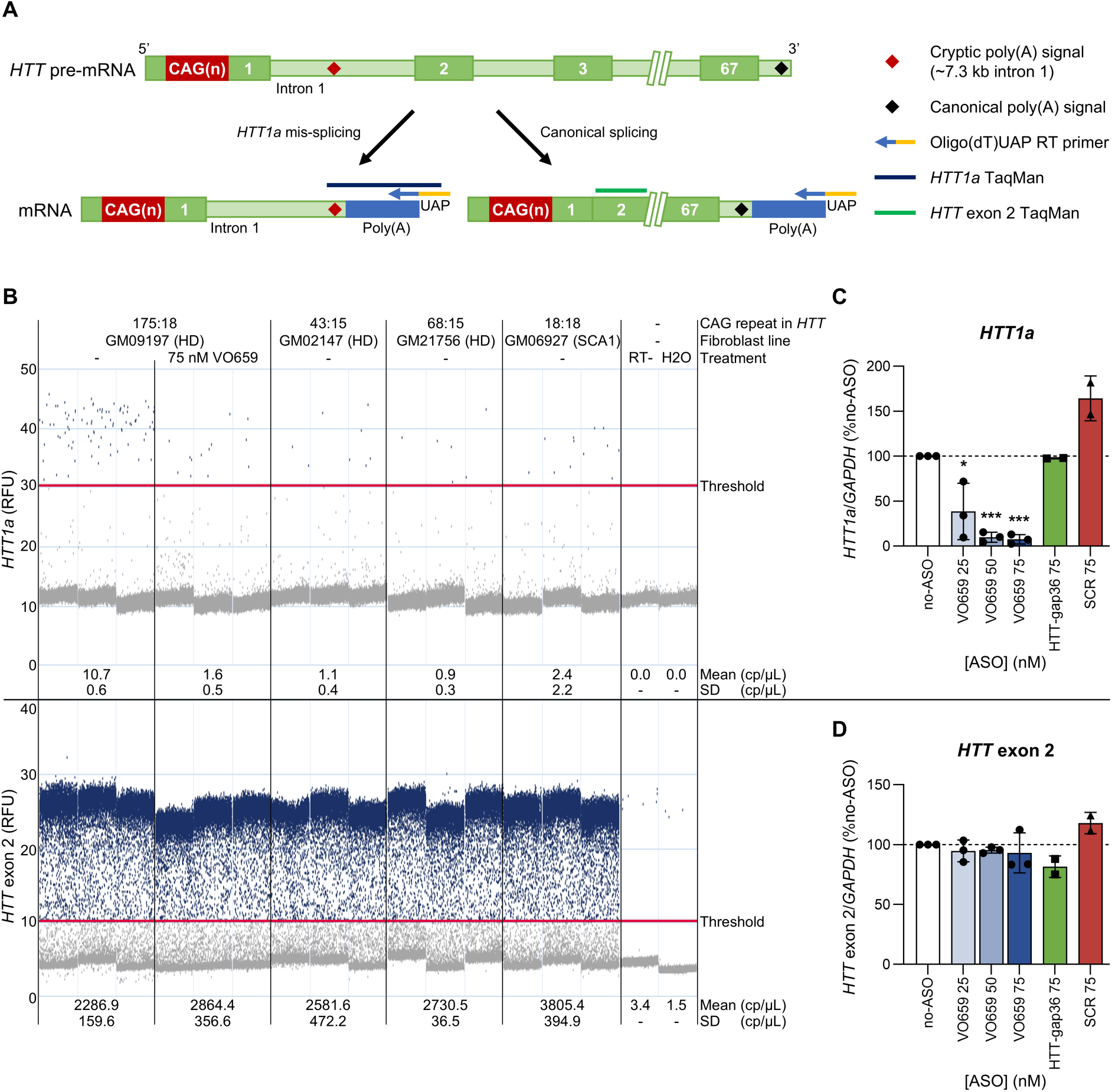
VO659 inhibits CAG repeat-mediated mis-splicing of HTT pre-mRNA that leads to *HTT1a formation* **A** Splicing schematic of *HTT* pre-mRNA, resulting in *HTT1a* and canonical *HTT* transcripts with corresponding polyadenine(poly(A)) tails. Indicated are the oligo(dT) primer with a universal adapter (UAP) sequence used for the reverse transcription (RT) reaction, and the *HTT1a* and canonical *HTT* exon 2 dPCR TaqMan assays as described by Hoschek et al..^26^ The reverse primer of the *HTT1a* taqman assay is complementary to the UAP overhang introduced during cDNA synthesis. **B** Representative RT-dPCR droplet images for the *HTT1a* (top) and *HTT* exon 2-containing transcripts (bottom) assays in various patient fibroblast lines with or without VO659 treatment, along with no-reverse transcriptase (RT-) and no-template (H2O) controls. Indicated concentrations represent the mean ± SD of 3 technical replicates. **C–D** *HTT1a* **(C)** and *HTT* exon 2-containing **(D)** transcript levels, corrected to a reference gene (GAPDH), in GM09197 HD patient fibroblasts (175:18 CAG) transfected with 25–75 nM ASO. Data are expressed as percentage of no-ASO control (set at 100%). The bars represent the average ± SD from 3 independent experiments. The data points represent the average per experiment of three technical replicates. Groups were compared to the no-ASO condition using a mixed-effects model (REML) with a post-hoc Dunnett’s test on non-normalized log-transformed data in GraphPad Prism. * = p<0.05, ** = p<0.005, *** = p<0.0005. RFU: relative fluorescent units; cp = copies

Transfection of 25–75 nM VO659 into the 175 HD patient fibroblast line led to a dose-dependent reduction of *HTT1a* mRNA expression, reducing it by ∼92% at the highest tested VO659 concentration (Fig. 4C) from ∼0.57% of total HTT mRNA levels to 0.04%. No effect on *HTT1a* levels was observed using 75 nM of the HTT-gap36 ASO, while SCR induced a 64% increase (Fig. 4C), possibly due to sequence-unrelated effects of transfection. VO659 did not affect *HTT* transcript levels containing exon 2 (Fig. 4D). As expected, HTT-gap36 did not reduce *HTT1a* mRNA levels as it targets *HTT* exon 36, although it did reduce *HTT* exon 2 levels by ∼18%, similar to the efficacy of HTT-gap36 on HTT protein levels observed in Fig. 2E. Transfection with 75 nM SCR resulted in an 18% increase of exon 2 containing HTT mRNA transcripts (Fig. 4D), similar to what was observed for the *HTT1a* transcript. Together, this shows that directly targeting the CAG repeat with VO659 can specifically inhibit mis-splicing of *HTT* pre-mRNA and thus prevent formation of the *HTT1a* transcript, in addition to sterically hindering its translation.

### VO659 inhibits the translation of toxic RAN proteins

RAN translation initiates within expanded repeats and generates short proteins from multiple reading frames that contain repetitive amino acid tracts that are aggregation-prone and have cytotoxic properties.^33,44^ Formation of RAN proteins is CAG repeat length dependent.^34^ We tested whether binding of VO659 to the expanded CAG repeat in *mHTT* mRNA could inhibit RAN translation. Unfortunately, we did not have access to antibodies recognizing endogenous polyA (GCA) or polyS (AGC) RAN proteins.^34^ We therefore made use of triple-tag overexpression constructs expressing *HTT* exon 1 with 20 or 101 CAG repeats with a different C-terminal tag (FLAG, HA or Myc) for each of the three different reading frames based on the design by Bañez-Coronel et al. (Fig. 5A).^34^ These constructs allow for quantification of the expression levels of polyQ, polyS, and polyA RAN proteins translated from the three reading frames, by detecting the FLAG, HA, or Myc tags, respectively. However, in our experiments, only the Myc-tagged polyA RAN proteins from reading frame 3 (RF3) could be consistently detected as a high molecular weight smear that was only present upon transfection of the 101 CAG, but not 20 CAG, repeat construct (Fig. 5B). Quantification of the Myc-tagged RF3 signal showed that VO659 dose-dependently reduced expression of these polyA RAN-proteins (Fig. 5C). At 35 and 75 nM, VO659 reduced expression of RF3-derived polyA proteins by 21 % and 56 %, respectively, while transfection with the SCR control did not affect polyA protein levels. In contrast, when detecting the canonical RF1 with a FLAG antibody, only a single band was observed at 32 kDa for the 20 CAG construct or at 158 kDa for the 101 CAG construct, corresponding to the full-length canonical open reading frame proteins (Supplemental Fig. 3A– B). Detection with an antibody targeting the N-terminus of HTT in a separate experiment showed a similar band pattern as the FLAG antibody, suggesting that most of the FLAG-tagged product also contained the HTT protein N-terminus and is therefore not RAN translation-derived (Supplemental Fig. 3C). We could not obtain a sufficiently reliable signal with the HA antibody to quantify RF2-derived HA-tagged polyS proteins. In some experiments, no clear HA signal was observed with both the 20 and 101 CAG constructs (Supplemental Fig. 3D), while in other experiments a faint smear of HA-tagged polyS RAN proteins was visible, with a higher-intensity signal for the 101 CAG construct than for the 20 CAG construct (Supplemental Fig. 3E). Treatment with VO659 did seemingly reduce the intensity of HA-tagged polyS signal when it was detectable. However, due to the low and inconsistent levels of polyS proteins from RF2, this could not be reliably quantified. These results are consistent with the literature, which has shown that translation from RF1 using constructs of this type predominantly produces the canonical ORF protein, that polyS proteins from RF2 are difficult to detect, possibly due to aggregation, and that polyA RAN translation from RF3 is the most robust.^45^ Similarly, in RAN translation constructs with the genomic context of spinocerebellar ataxia type 3 (SCA3), another CAG repeat-expansion disorder, polyS RAN translation was difficult to detect while polyA RAN translation was clearly observed.^46^

**Figure 5.**
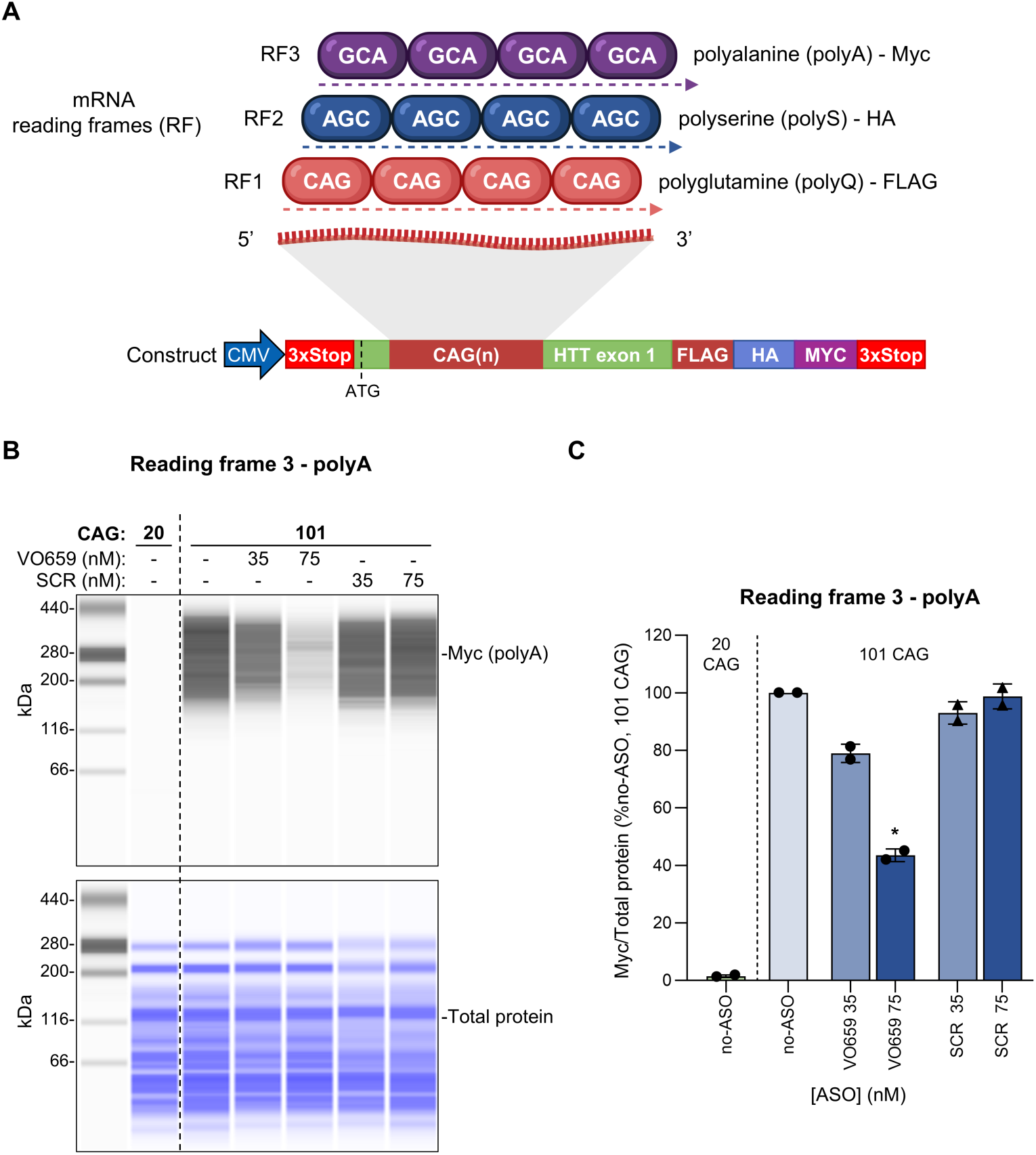
Translation of toxic RAN proteins is inhibited by VO659 **A** Schematic of out-of-frame translation from a *HTT* expression construct with 20 or 101 CAG repeats containing C-terminal tags in the three different reading frames. Translation from reading frame 1 (RF1), including canonical open reading frame (ORF) translation, results in expression of FLAG-tagged polyglutamine (polyQ) proteins, translation from RF2 results in expression of HA-tagged polyserine (polyS) proteins, and translation from RF3 results in Myc-tagged polyalanine (polyA) proteins. The construct contained a stop codon in each of the reading frames (3xStop). **B** Representative Simple Western (SW) blot views of HeLa cell protein lysate co-transfected with the construct described in A and 35–75 nM VO659 or scrambled (SCR), probed with an anti-Myc tag antibody (Cell Signaling Technology, #2278, 1:25) **C** Quantification with SW of Myc-tag (polyA) signal intensity in samples treated as described in B. Bars represent the average ± SD of the relative polyA RAN protein levels (No-ASO control of the 101 CAG condition set at 100%) of two independent experiments, corrected for total protein. Data points represent the average per experiment from 3 separate culture wells. Groups were compared to the no-ASO condition using a one-way repeated-measures ANOVA with a post-hoc Dunnett’s test on non-normalized log-transformed data in GraphPad Prism. * = p<0.05, ** = p<0.005, *** = p<0.0005

In conclusion, although RAN-translation could not be clearly detected in all three reading frames, the polyA data from RF3 suggests that binding of VO659 to an expanded 101 CAG repeat hinders translation of toxic RAN proteins.

### VO659 distributes well throughout the CNS *in vivo* with a long tissue half-life

Aside from the potential to reduce the various toxic protein isoforms derived from the expanded mutant *HTT* gene, it fundamentally important that therapeutics have a good pharmacokinetic (PK) profile and are sufficiently taken up in target tissues without being quickly metabolized or excreted. There are significant differences in uptake, stability and half-life between ASOs, largely determined by their chemical makeup and sequence. VO659 is a fully 2’-*O*-Me modified ASO with a PS backbone, which enhances nuclease resistance and stability and allows for good biodistribution and cellular uptake.^47^ Here, we compared the PK profiles of VO659 and HTT-gap36, a mixed PS/PO 2’-*O*-MOE modified gapmer. Wild-type C57BL/6J mice were given a single intracerebroventricular (ICV) equimolar dose of VO659, or HTT-gap36. The mice were sacrificed 1, 14, 35 or 84 days after dosing (Fig. 6A) and CNS and peripheral tissues (liver and kidney) were collected. The levels of ASO were determined using a sandwich hybridization ELISA. VO659 showed higher brain levels than HTT-gap36 at all time points across the collected CNS tissues. ASO levels were highest in every tissue 24 hours after dosing, after which a gradual decrease was observed. In cerebellum, the geometric mean (GM) level of VO659 was 16 µg/g on day 1, which decreased to 7 µg/g on day 84, corresponding to a tissue half-life of 69 days (Fig. 6B). HTT-gap36 levels were lower in the cerebellum, starting at 10 µg/g one day after dosing. On day 84, this decreased to 1 µg/g, with most samples measuring below the lower limit of quantification (LLOQ) of the ELISA assay, corresponding to a 2.5-fold shorter half-life of 27 days. In the other CNS tissues, a similar pattern was observed. The half-life of VO659 was 46, 36 and 26 days in hippocampus (Fig. 6C), cortex (Fig. 6D) and striatum (Fig. 6E), respectively, while HTT-gap36 showed 1.6 to 3.3-fold shorter half-lives of, respectively, 14, 22, and 10 days in the same tissues.

**Figure 6.**
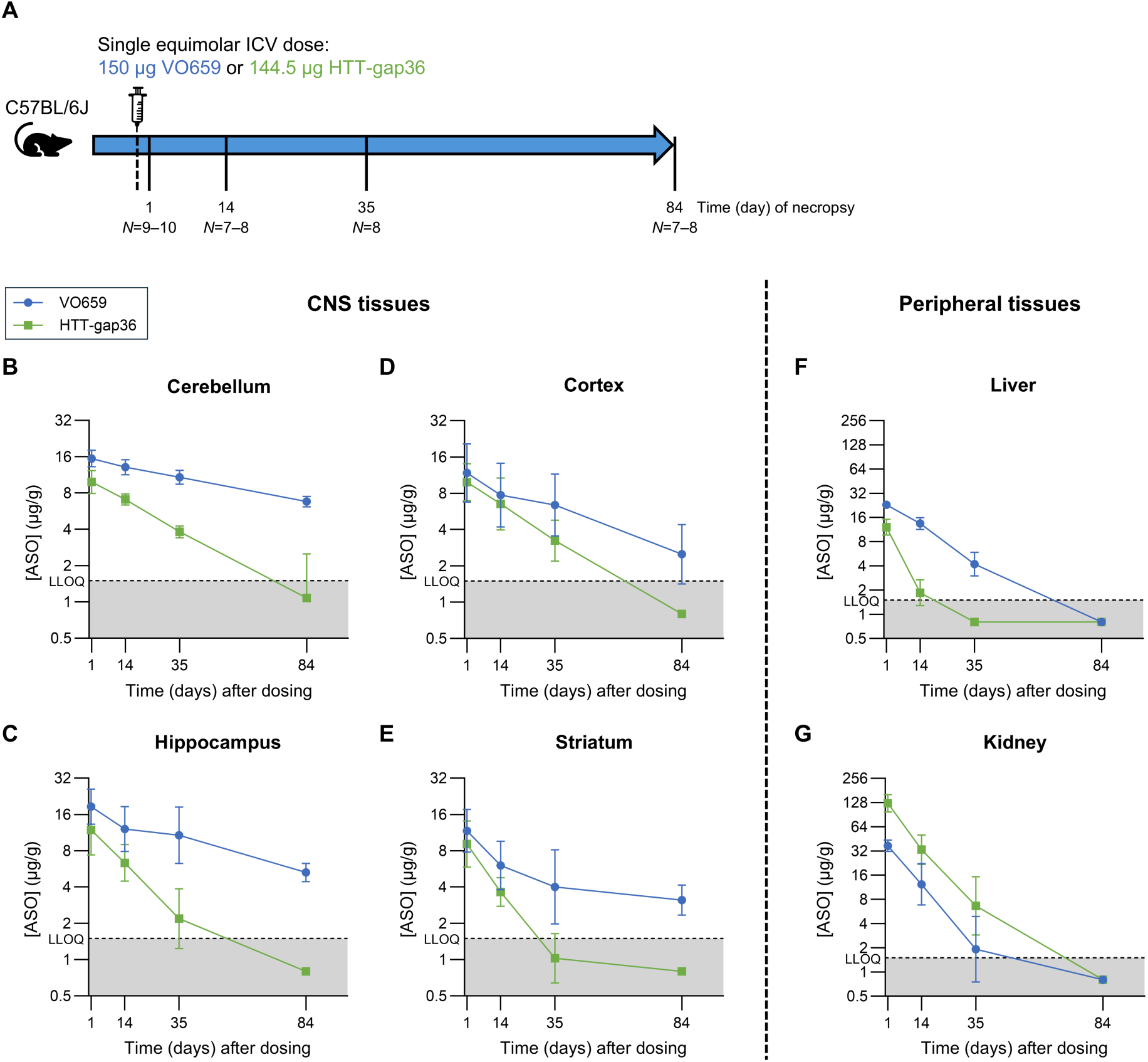
VO659 is taken up efficiently and has a long half-life throughout the CNS **A** Outline of an *in vivo* pharmacokinetic (PK) study in wild-type C57BL/6J mice. Mice were given a single intracerebroventricular (ICV) dose of VO659 or an equimolar equivalent of HTT-gap36 at 12 weeks of age. The mice were sacrificed at four time points as indicated and central nervous system (CNS), liver and kidney tissues were collected for PK analysis. The number of mice (*N*) per timepoint per ASO group is indicated. **B–G** Levels of ASO measured with a sandwich-hybridization ELISA. The data are expressed in µg ASO per gram of tissue (µg/g) in cerebellum **(B)**, hippocampus **(C)**, cortex **(D)**, striatum **(E)**, liver **(F)** and kidney **(G)**. The geometric mean (GM) ± geometric standard deviation (GSD) per timepoint is shown. The lower limit of quantification (LLOQ) of the ELISA assay was 1.5 µg/g, as indicated by the dashed lines. For plotting and calculation purposes, samples that measured below the LLOQ were given an arbitrary value of half of the LLOQ (0.75 µg/g).

While efficient CNS uptake is beneficial, minimizing ASO accumulation in liver and kidney is important from a safety perspective. One day after dosing, VO659 levels in liver were higher than HTT-gap36 at 23 µg/g and 12 µg/g, respectively. Although the half-life of VO659 in the CNS was long, its half-life in liver tissue was noticeably shorter at 15 days, with all liver samples showing levels below the LLOQ on day 84 (Fig. 6F). The half-life of HTT-gap36 in liver was 5 days. In kidney, HTT-gap36 levels were high on day one (126 µg/g), while VO659 levels were lower (37.1 µg/g). The half-life for both ASOs was comparably short in kidney, with 9 days for VO659 and 7 days for HTT-gap36 (Fig. 6G). Together, this shows that VO659 has a beneficial PK profile compared to HTT-gap36, with a higher CNS uptake and longer half-life in the mouse CNS after ICV dosing.

## Discussion

Despite the identification of the mutation causing HD over 30 years ago,^1^ there is still no approved disease-modifying therapy. Here, we show that the steric-blocking ASO VO659 has a unique HD-relevant MoA, reducing levels of multiple mHTT-derived toxic protein species that drive neurotoxicity in HD. VO659 is a fully modified 2’-*O*-methyl (2’-*O*-Me) phosphorothioate (PS) ASO that binds to the expanded disease-causing CAG repeat in exon 1 of *mHTT* (pre)mRNA and sterically hinders translation, leading to lower levels of mHTT protein while not affecting *mHTT* mRNA levels. VO659 has several advantages over other approaches, including allele-preferential reduction of mHTT protein and lowering of multiple toxic mHTT-derived protein species.

Using a combination of HD patient-derived fibroblasts and HeLa cells transfected with CAG repeat expression constructs, we confirmed that the efficacy of VO659 is CAG repeat length-dependent, leading to allele-preferential knockdown of mHTT protein levels with less reduction of wtHTT. The longer expanded CAG repeat in *mHTT* can accommodate more VO659 copies than the shorter repeat stretches in *wtHTT* transcripts, resulting in a stronger steric hindrance of translation. Although the function of wtHTT is not fully understood, it appears to be critical for normal brain function, playing an important role in several important cellular processes, including transcriptional regulation, vesicle and mitochondrial trafficking, transport of brain derived neurotrophic factor (BDNF), and synaptic transmission.^14^ Moreover, wtHTT may exert neuroprotective effects through anti-apoptotic activity.^48–49^ In adult mice, tamoxifen-induced conditional knockout of Htt resulted in progressive neurodegeneration, thalamic calcification, and behavioural and motor deficits.^50–51^ Furthermore, knockout of wtHtt in the YAC128 HD mouse model resulted in more severe motor dysfunction, hyperkinesia, testicular degeneration, impaired lifespan, and a modest worsening of striatal neuronal atrophy.^52^ However, partial reduction (up to 45%) of wtHTT in the putamen was well tolerated in rhesus macaques for up to 6 weeks,^53^ and heterozygous inactivation of HTT in humans with only one copy of *wtHTT* did not result in any phenotypic abnormalities or pathology.^54^ While this suggests that lowering wtHTT protein levels by up to 50% may be safe and well-tolerated in the absence of any mHTT, the long-term tolerability and the dominant-negative effects of mHTT’s presence remain unknown.

It is becoming increasingly clear that somatic expansion of the germline-inherited mutant CAG repeat is an important step in disease pathogenesis in HD. Genome-wide association studies (GWAS) have identified variants in multiple DNA mismatch repair genes, including *MSH3*, *MLH1*, *PMS1*, and *PMS2,* as genetic modifiers of disease age-of-onset in HD.^15,55–57^ These proteins play a role in repairing CAG extrusions *in vitro* and have been shown to modify somatic CAG-repeat instability *in vivo* in various mouse models.^58^ Somatic expansion of the CAG repeat in the *mHTT* gene results in widespread transcriptional dysregulation, ultimately resulting in neuronal death.^17^

We showed here that, although VO659 acts in both the nucleus and the cytoplasm, VO659 does not impact somatic expansion directly. However, it does effectively bind to transcripts with ultra-long CAG repeats that arise from somatic expansion. We demonstrated a clear CAG repeat length-dependent effect of VO659 in both the dual luciferase reporter assay and in HD patient fibroblasts. The highest efficacy of VO659 was observed in a fibroblast line derived from a juvenile HD patient with 175 CAG repeats. Through somatic instability, the mutant repeat can expand to beyond 400 CAGs.^17–18^ The efficacy and allele-preferential action of VO659 may even further increase with repeat lengths longer than 175 CAGs. However, this could not be tested directly, as it is difficult to clone constructs with these ultra-long (>150) CAG repeats and we had no access to cell lines with more than 175 CAGs in the *HTT* gene that are scarce with limited availability.

One of the pathways through which somatic expansion exerts neurotoxicity is the expression of the *HTT* exon 1-derived HTT1a protein.^22^ As shown by Hoschek et al.,^26^ longer CAG repeats result in more mis-splicing and higher expression of *HTT1a* mRNA. Consistent with this, we could only reliably detect *HTT1a* mRNA in the HD patient fibroblast line with 175 CAG repeats. There is increasing evidence that solely targeting full-length HTT without affecting HTT1a levels may not be sufficient to prevent HD pathogenesis.^59^ Indeed, in Q111 and zQ175 HD mouse models treated with ASOs or siRNAs, respectively, compounds that targeted HTT1a were clearly more effective at reducing HD-related aggregation and pathology than compounds that did not target HTT1a.^21,43^ Moreover, lowering the production of HTT1a by deleting the cryptic polyA sites from mouse Htt intron 1 in the HdhQ150 HD mouse model not only delayed the appearance of HTT1a aggregates by several months, but also prevented the disease-associated increase in CSF of neurofilament light chain (NfL), a sensitive biomarker of neuroaxonal damage that is increased in CSF of HD gene expanded individuals decades from clinical onset and predictive of imminent clinical decline.^22,60^ NfL is increasingly considered a biomarker relevant to interpret whether a HTT-lowering therapy is having positive or negative effects on the underlying disease pathway. Thus, while VO659 does not reduce somatic expansion directly, by lowering HTT1a levels, it reduces one of the proposed toxic mediators of somatic expansion.

Here, we showed that VO659 can act in two ways to keep HTT1a protein levels low: first, by preventing the mis-splicing leading to production of *HTT1a* transcripts, and second, by effectively inhibiting translation of any remaining *HTT1a* transcripts. The effect of VO659 on *HTT1a* mRNA levels is likely caused by sterically hindering splicing factors from aberrantly binding to the expanded CAG repeat in *mHTT* pre-mRNA, thus inhibiting activation of the cryptic polyadenylation site in *HTT* intron 1. In contrast, the HTT-gap36 gapmer that binds downstream of *HTT* exon 1 did not affect *HTT1a* mRNA levels, even though a small decrease of total *HTT* mRNA expression was observed. This corroborates the data from the previously mentioned Q111 and zQ175 HD mouse studies in which gapmers and siRNAs that bind downstream of *HTT* exon 1 did not affect *HTT1a* levels.^21,43^

We have previously shown that VO659 significantly reduced levels of both soluble and aggregated HTT protein in the brain of the R6/2 HD mouse model which expresses a human HTT exon 1 protein.^61^ However, we could not replicate this finding in the HD patient fibroblast line with a CAG repeat of 175, since we were unable to detect endogenous expression of HTT1a protein *in vitro* (data not shown), consistent with the very low *HTT1a* mRNA expression levels. Therefore, we used HTT1a overexpression constructs with 100 and 156 CAGs instead, showing that VO659 reduced HTT1a protein levels as well as the number of HTT1a aggregates in cells transfected with the constructs. Together, this showed that VO659 acts at the pre-mRNA level to block formation of the *HTT1a* transcript and, subsequently, at the mRNA level to block its translation to HTT1a protein.

Furthermore, mutant CAG repeats can lead to formation of secondary structures in *mHTT* mRNA that can result in the production of toxic RAN proteins from different reading frames.^31,34–35^ Besides RAN translation, translational frameshifting during translation of expanded CAG repeats in *mHTT* can also generate toxic polyA or polyS containing protein.^62–64^ In HD patients, it is uncertain whether RAN translation or frameshifting is the primary process that leads to expression of toxic frameshifted proteins, although these processes are not mutually exclusive. It is unclear whether the frameshifted proteins we observed using the triple-tag constructs were the result of RAN translation or ribosomal frameshifting, but for simplicity we refer to them here as RAN proteins. Regardless of the precise mechanism leading to expression of the frameshifted proteins, VO659 induced a dose-dependent reduction of levels of polyA frameshifted proteins detected by the anti-Myc antibody, illustrating that binding of VO659 to the expanded CAG repeat can block production of these toxic proteins, although this could not be confirmed for polyQ and polyS RAN proteins as these were not consistently detected.

RAN translation may also occur from mRNA transcribed from the antisense strand, *HTT-AS*.^34^ As this RNA contains an expanded CUG repeat rather than CAG repeat, it is unlikely that VO659 will reduce RAN translation from this transcript. However, the contribution of RAN translation of *HTT-AS* transcripts to HD pathogenesis is likely limited, as *HTT-AS* is generally expressed at low levels and is further reduced in the HD patient brain.^65^

The *mHTT* transcript itself may also contribute to disease onset and progression. In fact, blocking the activity of expanded CAG repeats in *mHTT* transcripts without modifying HTT RNA or protein levels was sufficient to reverse motor symptoms in the R6/2 HD mouse model.^66^ Secondary structure formation involving the elongated CAG repeat in *mHTT* mRNA, resulting in hairpins or stem-loop structures has been linked to RNA toxicity. ^31,67–69^ RNA toxicity is a well described pathogenic mechanism in several other (non-CAG) repeat expansion disorders like Myotonic Dystrophy type 1 and 2 (DM1/2) and C9orf72-related Amyotrophic Lateral Sclerosis (ALS).^70–72^ In these diseases, RNA secondary structures result in formation of RNA foci and sequestration of mRNA binding proteins leading to misprocessing of mRNA,^73–74^ a process that has also been observed in HD.^31,66,68,75^ We consider it likely that VO659 will also impact RNA toxicity, since binding of VO659 to the expanded CAG repeat could interfere with secondary structure formation and formation of RNA foci. However, we did not investigate this in the current study.

Besides having a MoA that targets multiple drivers of neurotoxicity, the uptake, biodistribution and half-life of VO659 are equally important. Because ASOs do not readily cross the blood-brain barrier (BBB), ASOs for neurodegenerative diseases must be injected directly into the cerebrospinal fluid (CSF) through intrathecal (IT) or intracerebroventricular (ICV) injections, after which they take advantage of the natural flow of CSF to distribute throughout the CNS and be taken up by neurons and other cell types.^76^ We showed that VO659, a fully PS 2’-*O*-Me modified ASO, has a more favorable PK profile compared to HTT-gap36, a mixed PS/PO 2’-*O*-MOE modified gapmer, with better uptake throughout the mouse CNS. Higher and more stable levels of VO659 than of HTT-gap36 were observed in all brain regions at all time points after ICV dosing. We also observed a long brain half-life in non-human primates (NHPs) (>150 days) (data not shown), similar to the long CSF half-life of steric blocking ASO Spinraza (163 days) which also has a full PS backbone.^77^ In line with this, an infrequent dosing regimen of once per 6 months is currently being explored in the ongoing VO659 phase1/2a clinical trial (NCT05822908). Interestingly, in peripheral tissues, the half-life of VO659 was short (1 to 2 weeks) similar, indicating that although VO659 is stable in CNS it does not accumulate in liver and kidney.

In patients, ASOs, including VO659, are often administered via an IT injection. A drawback of IT-delivered ASOs is their relatively poor distribution to deep brain, although it remains unknown to date how essential it is to reach deep brain regions for the treatment of HD. ASOs have a good uptake in the cortex, which is, besides the striatum, one of the most severely affected brain regions in HD, with the cortico-striatal synapse being one of the earliest and most affected sites in HD pathophysiology.^78–83^ BBB-shuttle technology, making use of the transferrin receptor (TfR1) to shuttle larger molecules into the CNS compartment, has been shown preclinically to achieve a more homogeneous ASO distribution across the entire brain including deep brain upon systemic delivery.^84–85^ This approach is currently being explored by Sarepta Therapeutics for HD in a recently initiated first-in-human phase 1 trial with SRP-1005, an investigational HTT-lowering siRNA coupled to a monovalent fragment antigen binding (fAb) of TfR1 and is dosed using subcutaneous injection (NCT07536061). This approach may lead to a better benefit/risk of the ASO approach in HD via improved brain distribution.

While there is no approved disease modifying treatment yet available, it is an exciting time for the HD field given the wide diversity of HTT-lowering modalities in development aimed at halting or slowing down disease progression. These modalities include ASOs, siRNAs, gene therapy and small molecules, and differ fundamentally in whether they are pan-HTT-lowering or allele-preferentially lower mHTT. Moreover, they differ in whether they exclusively lower full-length mHTT or also HTT1a protein levels and differ in their dosing frequency and route. Most HTT-lowering therapeutic approaches currently in clinical development for HD are pan-HTT-lowering and do not target HTT1a. VO659 is the only approach currently in clinical development that is both allele-preferential and lowers HTT1a.

An allele-preferential HTT-lowering therapeutic approach offers clear advantages given the important implicit role of wtHTT in maintaining cell function and homeostasis. As suggested by the clinical results of tominersen (non-allele-preferential, non-*HTT* exon 1 targeting ASO), too much lowering of wtHTT protein, particularly in more clinically progressed patients at baseline, may lead to negative clinical outcomes.^37^ However, it is more challenging to develop an allele-preferential therapeutic approach given the high similarity between *wtHTT* and *mHTT*. Wave Life Science’s WVE-003 ASO (NCT05032196) and Roche’s recently discontinued ASO RG6496 (NCT07246941) both target a single nucleotide polymorphism (SNP) variant in *HTT* that is frequent on HD chromosomes, allowing allele-preferential reduction of mHTT protein while having only a limited effect on wtHTT protein. A limitation of SNP targeting approaches is that they are only applicable in a subset of HD patients that carry the SNP variant on the *mHTT* allele, e.g. up to 40% of the HD population for the WVE-003 targeting *rs362273* (SNP3).^86^ In contrast, VO659 is applicable in the entire HD population, and, additionally has therapeutic potential for all nine CAG repeat-expansion disorders, which, besides HD, includes spinocerebellar ataxia types 1, 2, 3, 6, 7, and 17 (SCA1, 2, 3, 6, 7, 17), dentatorubral-pallidoluysian atrophy (DRPLA) and spinal and bulbar muscular atrophy (SBMA).^87^ We previously showed that VO659 reduced mutant ataxin-1 and ataxin-3 protein levels in patient fibroblasts as well as in SCA1^154Q/2Q^ and SCA3 MJD84.2 mouse models, respectively.^88^

Only a few therapeutic approaches have the potential to reduce HTT1a expression by directly targeting *HTT* exon 1. The only two candidates in clinical development which target HTT1a, besides VO659, are ALN-HTT02 and AMT-130, which are both non-allele-preferential, equally lowering wtHTT and mHTT protein levels. ALN-HTT02, in development by Alnylam Pharmaceuticals, is an IT-delivered siRNA therapy that non-allele-preferentially lowers *HTT* mRNA levels and targets *HTT* exon 1 and is currently being tested in a phase 1 clinical trial to assess safety (NCT06585449). AMT-130 is a non-allele-preferential AAV5 gene therapy in development by uniQure, delivering a miRNA targeting HTT exon 1. AMT-130 is a one-time intracranially administered gene therapy that requires stereotactic intracerebral infusion directly into the caudate and putamen (NCT04120493).

There are two small molecules in clinical development for HD, votoplam (PTC-Novartis; NCT07326709) and SKY-0515 (Skyhawk Therapeutics; NCT06873334). The main advantage of these small molecules is that they can cross the blood-brain barrier, allowing for oral dosing, and in principle have a more uniform brain distribution. Both votoplam and SKY-0515 non-allele-preferentially reduce HTT levels by inducing inclusion of a pseudo exon into mature *HTT* mRNA leading to nonsense-mediated decay. Interestingly, SKY-0515 also reduces levels of PMS1, which, as mentioned before, is a modifier of somatic expansion of the CAG repeat and therefore has the potential to directly slow somatic expansion.^89^

Compared to the HTT-lowering therapies outlined above, VO659 has a unique mode of action. VO659 is the only therapy currently in clinical development for HD that potentially addresses all the CAG repeat-related toxic mechanisms of actions, that include expression of the full-length mutant HTT protein, expression of the toxic HTT1a isoform, and CAG repeat secondary structure-related toxicities. Furthermore, due to its CAG repeat-dependent efficacy, VO659 allele-preferentially reduces mutant HTT protein levels, becomes more potent as somatic expansion of the CAG repeat occurs and has potential for the entire class of CAG repeat expansion disorders. Furthermore, we showed that VO659 has a beneficial pharmacokinetic profile and a long half-life in the CNS, potentially allowing for infrequent dosing of patients. VO659 is currently being tested in a phase 1/2a clinical basket-case trial with HD, SCA1 and SCA3 patients in a once per 6 months dosing regimen to examine its safety profile in humans and look at early markers of target engagement. Future clinical data will inform on whether the unique promise of VO659’s mechanism as outlined in this manuscript will translate to the clinic.

## Materials and Methods

### Cell culture and seeding

Patient and control fibroblast cell lines were obtained from the Coriell Institute for Medical Research (Camden, NJ). HeLa CCL-2 cells were obtained from the American Type Culture Collection (ATCC, Manassas, VA). The length of the CAG repeats in HTT in the fibroblast lines was determined with Sanger sequencing (Table 1). Fibroblasts were cultured in Minimum Essential medium (MEM) (Gibco, Thermo Fisher Scientific, Waltham, MA, #10370070) with 15% fetal bovine serum (FBS) (Gibco, #A5670701), 1% penicillin/streptomycin (Gibco, #15140122) and 1% GlutaMAX supplement (Gibco, #35050038). HeLa cells were cultured with 10% FBS, but otherwise the same medium. All cells were maintained in a 37 °C incubator with 5% CO2.

**Table 1.** Overview of used cell lines.

| Cell line | Supplier | polyQ disease | CAG length in *HTT* ** |  |
| --- | --- | --- | --- | --- |
|  |  |  | Allele 1 | Allele 2 |
| GM09197* | Coriell | HD | 175 | 18 |
| GM21756* | Coriell | HD | 68 | 15 |
| GM02151 | Coriell | HD | 45 | 18 |
| GM00305 | Coriell | HD | 44 | 17 |
| GM02147* | Coriell | HD | 43 | 15 |
| GM06927 | Coriell | SCA1 | 18 | 18 |
| GM02171* | Coriell | - | 17 | 15 |
| HeLa | ATCC | - | NR | NR |

\*CAG repeat lengths previously reported in Röttgering et al.<sup>38</sup> \*\*CAG repeat length determined with Sanger sequencing defined as the stretch of pure CAGs before any CAA interruptions at the 3' end of the repeat in *HTT*. NR = Not reported

For passaging, cells were washed with phosphate-buffered saline (PBS) (Gibco, #14190144), detached with 0.05% trypsin-Ethylenediaminetetraacetic acid (EDTA) (Gibco, #25300054) for 5 minutes (min) at 37 °C, and plated on new culture dishes with fresh culture medium. When seeding of a specific number of cells was required, viable cells were counted using 0.4% trypan blue with a Luna-II automated cell counter (Logos Biosystems, Gunpo-si, South Korea, #L40002).

### ASO transfection of fibroblasts

To transfect fibroblasts with ASOs, fibroblasts were seeded on 10-cm dishes in 6 mL culture medium. Because the fibroblast lines grew at different rates, the number of cells seeded and the time between seeding and transfection varied between cell lines, ranging from 250,000– 612,000 cells per 10-cm dish, and from 24–96 h before transfection. To ensure comparable transfection conditions across the various fibroblast lines, cells were only transfected once the 10-cm dishes reached 70-80% confluency. On the day of transfection, the 10-cm dishes were refreshed with 6 mL transfection medium containing 10% instead of 15% FBS. Then, ASOs were transfected using Lipofectamine RNAiMAX (Invitrogen, Thermo Fisher Scientific, Waltham, MA, #13778150) according to the manufacturer’s protocol. Lipofectamine RNAiMAX and ASOs were separately diluted in Opti-MEM (Gibco, #31985062). The dilutions were mixed 1:1 (v/v) and incubated for 5 min at room temperature (RT) before 1.5 mL of the ASO-lipid complexes was added to the 10-cm dishes, resulting in final ASO concentrations of 25–100 nM. The amount of Lipofectamine RNAiMAX per 10 cm dish was 3–12 µL, scaling with the amount of ASO transfected. For the non-treated (NT) control, 1.5 mL of Opti-MEM without ASO or transfection reagent was added. The fibroblasts were cultured for 96h in the presence of the transfection mix before harvesting. All ASOs used in this manuscript are listed in Supplemental Table S1 and were synthesized in house (VICO Therapeutics, Leiden, the Netherlands).

### ASO and plasmid co-transfection of HeLa cells

Plasmids were generated in-house using different backbones depending on the construct. Dual luciferase reporter (DLR) constructs were generated using a backbone obtained from Promega (Madison, WI, #C8021), while HTT1a overexpression constructs were generated using a custom pRP backbone obtained from VectorBuilder (Chicago, IL). The RAN translation constructs were fully synthesized by VectorBuilder. Schematics of all plasmids can be found in Supplemental Fig. S4A–C. ASOs used in co-transfection experiments can be found in Supplemental Table S1.

For co-transfection experiments with plasmids and ASOs, HeLa cells were used in different plate formats (6-well or 96-well), with specific seeding densities and transfection conditions depending on the assay (Supplemental Fig. S4D–E). In short, HeLa cells were seeded in culture medium and incubated at 37 °C for 24–48 h. Then, the medium was refreshed and the first transfection performed. After 4 h, medium was refreshed again, followed by the second transfection. Whether ASO or plasmid was transfected first was assay dependent (Supplemental Fig. S4E). Plasmids were transfected using either Lipofectamine-2000 (Invitrogen, #11668019) or Lipofectamine-3000 (Invitrogen, #L3000001), whereas ASOs were transfected with Lipofectamine RNAiMAX or Lipofectamine-2000. All transfection complexes were prepared in Opti-MEM according to the manufacturer’s instructions. Additionally, for plasmid transfections using Lipofectamine-3000, 0.15 µL of P3000 reagent was added per well of a 96-well plate. Specific transfection reagents, payload amounts (ng plasmid or nM ASO), and culture volumes for each assay are detailed in Supplemental Fig. S4E. Cells were harvested or fixed 24–72 h after the second transfection.

### Simple Western capillary immunoassays

Protein was isolated from cells using CelLytic M (Sigma-Aldrich, St. Louis, MO, #C2978) supplemented with cOmplete protease inhibitor (Roche, Basel, Switzerland, #04693116001), 1 tablet per 10 mL. Cells were trypsinized, spun down (350 g for 7 min), and lysed for >20 min at 4 °C with vortexing every 2-3 min. After lysis, samples were spun down (18000 g for 10 min) and supernatant was collected. Protein concentration was determined using the Pierce BCA kit (Thermo Fischer Scientific, #23227).

Simple Western (SW) capillary immunoassays were performed with the Wes (Bio-Techne, Minneapolis, MN, #004-600), Jess (Bio-Techne, #004-650) or Abby (Bio-Techne, #004-680) systems according to the manufacturer’s instructions. For all assays, 12–230 kDa (Bio-Techne, #SM-W001) or 66–440 kDa (Bio-Techne, #SM-W005) SW modules were used. Briefly, protein lysates were diluted to the desired concentration in 0.1X Sample Buffer and Fluorescent Master Mix, denatured at 95 °C for 5 min, and loaded into SW assay plates together with primary antibody (for antibody details and loading concentrations see Supplemental Table S2), HRP-conjugated anti-rabbit secondary antibody (Bio-Techne, #042-206), and luminol/peroxide detection reagent (Bio-Techne, #DM-003). For normalization, protein lysate was stained for total protein using the Total Protein Detection Module (Bio-Techne, #DM-TP01). When using the Wes system, total protein was detected on a separate plate using a protein aliquot from the same sample dilution as used for the antibody plate. When using the Jess or Abby systems, total protein was detected on the same plate, in the same capillaries as the antibody stain, using the RePlex Module (Bio-Techne, #RP-001). For the analysis of frame-shifted proteins Replex runs with 2 consecutive antibodies (anti-Myc or anti-HA followed by anti-FLAG, Supplemental Table S2) were also performed, and the resulting values were normalized using the average total protein value obtained from other plates. For every SW assay, the default parameters (separation time, separation voltage, antibody incubation time, and exposure) were used. The SW runs were analyzed with the Compass for SW software (Bio-Techne).

### Caspase-3 based HTT SW capillary immunoassay

WtHTT and mHTT protein levels were quantified using SW capillary immunoassays. However, because wtHTT and mHTT proteins do not readily size-separate on the SW platform, protein lysate was treated with caspase-3 before loading on SW assay plates, as we have previously described.^38^ Caspase-3 cleavage of HTT generates N-terminal HTT fragments that contain the polyQ region. In these N-terminal fragments, the polyQ region makes up a relatively larger portion of the protein, allowing for size-separation. In short, 0.26 mg/mL of protein lysate was treated overnight at 37 °C with 10.5 U/µL caspase-3 (Sigma-Aldrich, #235417) in a 10 µL reaction volume in the manufacturer-recommended buffer (0.1% CHAPS (m/v), 10% Glycerol (v/v), 100 mM NaCl, 10 mM DTT, 1 mM EDTA, 50 mM HEPES, pH 7.4). After incubation, caspase-3 treated samples were loaded on SW assay plates at a final concentration of 0.07 mg/mL and HTT and total protein were detected as described earlier.

### HTT dual luciferase reporter assay

Dual Luciferase Reporter (DLR) constructs were generated in house using the psiCHEK-2 backbone (Promega, #C8021). The constructs contained the coding sequence of *HTT* exon 1–6 (with 5–98 CAG repeats) fused upstream to a renilla luciferase (hRluc) via a flexible linker, and Firefly (Fluc) luciferase internal control under a separate promoter. HeLa cells were seeded in white clear-bottom 96-well plates (Greiner Bio-One, Kremsmünster, Austria, #655083) and transfected with 0.5–60 nM VO659, HTT-gap36 or SCR and the DLR constructs as described in the co-transfection section (Supplemental Fig. S4). Both HTT-hRluc fusion and Fluc reporter activities were measured 24h after transfection using the Promega Dual-Glo Luciferase assay system (Promega, #E2920) according to manufacturer instructions. In short, Fluc activity was measured by adding 75 µL Dual-Glo luciferase assay reagent per well to the transfection medium. After 15 min of shaking at 600 rpm (Heidolph, Schwabach, Germany, # 544-11200-00), luminescence was measured in a Synergy H1 plate reader (BioTek Instruments, Winooski, VT). Then, HTT-hRluc reporter activity was measured by adding 75 µl Dual-Glo Stop & Glo reagent to each well, followed by 15 min shaking at 600 rpm and luminescence readout. To determine ASO efficacy, the HTT-hRluc/Fluc ratio was calculated to correct for construct transfection efficiency and cell density. Then, the HTT-hRluc/Fluc ratio was expressed as percentage of untreated control, per construct.

### HTT1a immunostaining and imaging

HTT1a overexpression constructs with 100 or 156 CAG repeats under control of a cytomegalovirus (CMV) promoter and a GFP internal control under human phosphoglycerate kinase (hPGK) promoter were generated in house, using a custom pRP backbone obtained from VectorBuilder. As described in the co-transfection section and supplemental Fig. S4, HeLa cells were first transfected with 5–50 nM of VO659, HTT-gap36 or SCR, followed by transfection with 60 ng of the HTT1a overexpression constructs in black clear-bottom 96-well plates (Corning, #CLS3603). Three wells (technical replicates) were transfected per experiment. Then, 72 hours after transfection, the cells were washed twice with 100 µL PBS, followed by fixation with 50 µL 4% formaldehyde solution (Invitrogen, FB002) for 15 min at RT. After fixation, cells were washed three times with PBS and stored in PBS at 4 °C or immediately stained for HTT1a. For staining, cells were permeabilized with 100 µL 0.2% Triton X-100 in PBS for 20 min at RT, washed twice with PBS, and blocked with a 5% FBS and 0.5% Tween-20 in PBS solution for 45 min at RT. HTT1a was probed with the rabbit anti-HTT1a NEO p90 antibody, clone 11G2 (CHDI Foundation, New York City, NY, #CHDI-90004291),^41^ diluted 1:500 in blocking buffer, overnight at 4 °C. Then, following three washes with PBS, cells were incubated for 2 h with 1:1000 goat anti-rabbit Alexa Fluor 647 (Invitrogen, #A-21244) and a DAPI nuclear counterstain (1:1000) in blocking buffer. After three final washes with PBS, cells were imaged for visualization with a laser scanning confocal microscope (Carl Zeiss AG, Oberkochen, Germany, #LSM 800) using a 20x lens and the Zen Blue software v2.6 (Carl Zeiss AG).

For quantification of fluorescence intensities, plates were scanned using a CellInsight CX7 LZR high content imager (Thermo Fisher Scientific, #CX7A1110LZR) with a 20x lens using the wide field imaging option. A minimum of 20 fields were imaged per well for DAPI (405 nm channel), GFP (488 nm channel), and HTT1a (647 nm channel). Fluorescence intensities were quantified using a colocalization protocol from the HCS Studio v6.6.4 Cellomics Scan Software (Thermo Fischer Scientific). To summarize fluorescence intensity data, the median single-cell HTT1a intensity (corrected for GFP intensity) was calculated per technical replicate (e.g. culture well). These medians were then averaged to calculate a mean for each experimental condition. Additionally, high-intensity HTT1a foci (aggregates) were specifically detected by setting an intensity threshold and applying filtering based on size and shape such that diffuse HTT1a signal was excluded. The percentage of GFP positive cells that contained at least one HTT1a foci was then calculated per condition.

### HTT1a RT-dPCR

To quantify *HTT1a* and canonical *HTT* mRNA levels, HD patient fibroblasts were trypsinized and lysed in 500 µL NucleoZol (Machery-Nagel, Düren, Germany, #740404). RNA was isolated by adding 200 µL H_2_O to the lysed samples and incubating at RT for 5 min. After centrifugation at 18,000 g for 15 min, the upper aqueous phase was collected. RNA was precipitated by adding an equal volume of isopropanol for ∼15 min at RT and pelleted by centrifuging at 18,000 g for 15 min. The resulting RNA pellet was washed three times with 500 µL 80% ethanol and dissolved in H_2_O. RNA concentration was determined using NanoDrop (Thermo Fischer Scientific, #ND-ONE-W).

For cDNA synthesis, 2 µg of RNA was reverse transcribed using SuperScript IV reverse transcriptase (Invitrogen, #18090050) according to the manufacturer’s specifications, in a reaction volume of 40 µL. The reaction contained 1 µM of an oligo(dT) primer with a universal adapter sequence (RT-oligo(dT)UAP) based on a previously published protocol by Hoschek et al.^26^ (Supplemental Table S3). To control for gDNA contamination, a control without reverse transcriptase (RT-) was included in every experiment.

*HTT1a* and *HTT* canonical mRNA levels were quantified using multiplexed TaqMan assays and digital PCR (dPCR), using assays designed by Hoschek et al..^26^ In short, 12 µL dPCR duplex reactions were prepared using the QIAcuity Probe PCR Kit (QIAGEN, Venlo, The Netherlands #250101) according to manufacturer’s specifications. Each duplex reaction contained 4 µL of undiluted cDNA, 3 µL of 4X QIAcuity Probe PCR Master Mix, and both the *HTT1a* (FAM) and *HTT* canonical (HEX) assays at final concentrations of 263 nM for each primer (forward and reverse) and 138 nM for each TaqMan probe. Reactions were loaded on QIAcuity Nanoplate 8.5K 96-well plates (QIAGEN, #250021), which were sealed using Nanoplate seals (QIAGEN, #250099). The dPCR was performed on the QIAcuity eight (QIAGEN, #911050) system using standard priming and imaging settings, and a 2-step PCR protocol (initial denaturation of 2 min at 95 °C, then 40 cycles of 15 sec at 95 °C and 60 sec at 58 °C). For normalization, GAPDH reference gene levels were measured separately with the same protocol using a premade singleplex TaqMan assay (Thermo Fisher Scientific, #Hs04420632_g1, FAM-MGB) at a 1:60 (v/v) concentration and 4 µL of 1:500 (v/v) diluted cDNA. RT- and no template (H_2_O) controls were included in every dPCR run. Data were analyzed using the QIAcuity Software Suite (QIAGEN). Thresholds were set manually per TaqMan assay and applied uniformly across all wells within a plate run. All primer and probe sequences are listed in Supplemental Table S3.

### ASO dosing of Q111 mice

We used heterozygous Q111 mice (RRID: IMSR_JAX: 003598) on a C57BL6/J background bred at the University of Washington.^40^ Mice were CAG-sized with an average CAG length of 114 ± 3. Mice were fed ad libitum and maintained with a 12-hour light/12-hour dark cycle. Unilateral ICV bolus injections, which result in efficient brain-wide distribution of ASO, were conducted as previously described.^90^ Briefly, mice were anesthetized with isoflurane, placed on a stereotaxic device, and unilaterally injected with 200 μg of ASO in 10 μl of artificial cerebral spinal fluid (aCSF; Tocris Bioscience, Bristol, UK, #3525) at 2.5 months of age. All animal work was approved by the University of Washington animal use committee (protocol 4563-02).

### MSD quantification of wtHTT and mHTT protein levels

For MSD assays, cortical lysates were prepared in a non-denaturing lysate buffer and read on a MESO QuickPlex SQ120MM plate reader (Meso Scale Discovery) as previously described.^51^ The capture and sulfo-tagged detection antibody pairs were as follows. WT HTT: CHDI-90002133 (Coriell), D7F7 (CST); mHTT: 2B7 (Coriell), MW1 (DSHB).

### Somatic expansion analyses

Genomic DNA from Q111 mice (RRID:IMSR_JAX:003598) was extracted from striatal tissue, fluorescent CAG repeat-spanning amplicons were produced by PCR and subsequently measured on a Genetic Analyzer (Applied Biosystems) as previously described^39^ with the following primer pair: CAG1 (5’ ATGAAGGCCTTCGAGTCCCTCAAGTCCTTC 3’) and reverse primer Hu3 (5’ GGCGGCTGAGGAAGCTGAGGA 3’). PCR product size distributions were generated using Peak Scanner 2 software (Thermo Fisher Scientific) and subsequently imported into R for calculation of expansion indices according to previously described methods.^39,91^

### ASO dosing of C57BL/6J mice for PK analysis

Animal experiments (Study number C0250420) were carried out at Charles River Laboratories (Kuopio, Finland; accredited by AAALAC International) as specified in the license authorized by the national Animal Experiment Board of Finland and according to the National Institutes of Health (Bethesda, MD, USA) guidelines for the care and use of laboratory animals.

A total of 68 C57BL/6J mice (34 males, 34 females) were randomly assigned to the different treatment groups, maintaining an equal gender ratio. Male and female mice were stratified by weight to ensure similar body weights across all groups and timepoints. Mice were injected intracerebroventricularly (ICV) at 12 weeks of age with 150 µg VO659 (N=34) or an equimolar amount (144.5 µg) of HTT-gap36 (N=34), formulated in aCSF (Tocris Bioscience, #3525). Before injection, mice were anesthetized using isoflurane (2.0–5.0% induction, 1.0–2.5% maintenance) delivered in 70% N_2_O and 30% O_2_ at a flow rate of 600 mL/min. The ASO injections were given in a 5 µL volume at a rate of 0.6 µL/min with a stereotactically guided Hamilton syringe and infusion system (Harvard Apparatus, Holliston, MA) in the right lateral ventricle at Bregma coordinates AP: −0.5 mm, ML: +1.0 mm and DV: −2.5 mm. After injection, the needle was left in place for 5 min before withdrawal. Two to three mice were housed per cage and had *ad libitum* access to standard chow and water. Mice were euthanized at 24 h (*N*=9–10), 14 days (*N*=7–8), 35 days (*N*=8), or 84 days (*N*=7–8) after dosing with a pentobarbital (180 mg/kg) intraperitoneal injection, followed by brain isolation and collection of cortex, striatum, hippocampus and cerebellum. Additionally, one liver lobe and the left kidney were collected. Tissues were snap frozen in isopentane in a bath of ethanol on dry ice and stored at −80 °C until shipment to VICO Therapeutics, Leiden, the Netherlands. Storage at VICO was continued at −80 °C until PK analysis.

### Sandwich hybridization ELISA

To quantify ASO levels in tissue samples, a weighed amount of tissue was added to tubes containing Lysing Matrix D (MP Biomedicals, Santa Ana, CA, #1169130-CF), followed by 1 mL per 60 mg tissue of 2 mg/mL proteinase K (Invitrogen, #25530-031) diluted in a buffer containing 100 mM Tris, 200 mM NaCl, 0.2% (m/v) sodium dodecyl sulfate (SDS) and 5 mM EDTA. The tubes were placed in a MP FastPrep-24 5G Tissue lyser (MP Biomedicals, Santa Ana, CA, #116005500) and homogenized for 4×20 sec at 10 m/s, with cooling on ice after two rounds. After homogenization, CNS and liver tissue samples were incubated for 2 h, while kidney samples were incubated for 4 h, at 55 °C under gentle agitation.

ASO levels in tissues were detected using a full-length product (FLP) specific sandwich hybridization enzyme-linked immunosorbent assay (ELISA). Eight standard curve calibrators (range 1.5–40.0 ng/mL) of the appropriate reference ASO were prepared through serial dilution in 1 mg/mL untreated control tissue lysate. Tissue samples were diluted 1:60 (v/v) in PBS to a tissue concentration of 1 mg/mL and further diluted to concentrations within the calibration curve range using 1 mg/mL untreated control lysate. This approach was used to account for possible matrix effects by keeping tissue concentrations the same between calibrators and study tissue samples. For CNS sample analysis, a pool of 1 mg/mL untreated control CNS tissue lysate was used. For liver and kidney analysis, a 1:1 (v/v) pooled mixture of 1 mg/mL untreated liver and kidney control lysate was used.

Streptavidin coated 96-well plates (Roche, #11734776001) were incubated for 30 min at 37 °C with 25 nM of a biotinylated capture probe with a reverse-complement sequence for either VO659 or HTT-gap36 (Supplemental Table S4). After 4x washing with Tris-buffered saline with Tween-20 (TBS-T, Sigma-Aldrich, #91414), the tissue samples, calibrators, and an untreated control sample were incubated on the plate *in duplo* for 30 min at 37 °C. Then, plates were washed 4x with TBS-T and incubated with 2 nM of a digoxigenin (DIG)-labelled detection probe which was reverse complementary to the 5’ overhang of VO659 or HTT-gap36 that was not directly bound to the capture probe (Supplemental Table S4). The plates were incubated for 30 min at 45 °C for VO659 and at 37 °C for HTT-gap36. Capture and detection probes were diluted in a buffer containing 58 mM Na_2_HPO_4_, 917 mM NaCl, and 0.24% Tween-20 (v/v). After 4 washes with TBS-T, the DIG label was probed with 1:7500 (v/v) anti-DIG horse-radish peroxidase antibody (Roche, #11207733910) in 1% milk (m/v) in TBS-T for 30 min at RT. Then, the plates were washed 4 times with TBS-T, and incubated with 3,3’,5,5’-Tetramethylbenzidine (TMB) (Sigma-Aldrich, #T4444) for 15-20 min at RT, after which the reaction was stopped using 345 mM maleic acid. Absorption was measured at 450 nm in a Synergy HT plate reader (BioTek Instruments). Raw absorption data was processed using the Gen5 software v1.09 (BioTek Instruments). A four-parameter logistic regression model standard curve was fitted to the background corrected calibrator data to estimate ASO concentrations in study tissue samples. Final ASO concentrations are reported as µg ASO per gram of tissue (µg/g). Because the minimum tissue dilution was 1:60 (v/v), the lower limit of quantification of the ELISA assay was 1.5 µg/g. All samples that measured below the LLOQ, were given an arbitrary value of 0.75 µg/g, half of the LLOQ, for half-life calculations and plotting purposes. For all PK data, the geometric mean (GM) with the geometric standard deviation (GSD) was calculated and plotted. ASO half-lives were determined based on a nonlinear curve fit between the maximum concentration (Cmax) and the end concentration (last timepoint). In cases where multiple timepoints measured below the LLOQ, the first timepoint with a GM concentration below the LLOQ was used as an endpoint.

### Software and statistics

All data graphs were created and all statistics were performed with GraphPad Prism v10.2.3 (GraphPad Software, Boston, MA). The illustrations in graphical abstract/Fig. 1 (https://BioRender.com/xzr5h8w) and Fig. 5A/Fig. S3A (https://BioRender.com/t5mhcii) were created using BioRender. Somatic expansion indices were calculated using R.

## Data availability statement

The data analyzed in this study may be obtained from the corresponding author upon reasonable request.

## Acknowledgments

We thank Prof. Dr. Willeke van Roon-Mom from (Leiden University Medical Center), Scott Schobel, MD (VICO Therapeutics BV) and Dr. Prarthana Khanna (VICO Therapeutic BV) for critical review of the manuscript. This work was funded by VICO Therapeutics BV.

## Author contributions

B.R. and N.D. drafted and revised the manuscript. B.R., R.V., S.B. and C.B., visualized the data. R.V. made the graphical abstract. J.F.C and R.M.B performed and analyzed the Q111 mouse experiments and wrote the corresponding material and methods sections. B.R., R.W., S.B., R.V., J.T., C.B., developed methodologies, performed experiments and analyzed data for the other figures in the manuscript. L.T., J.T., and N.D. supervised experimental work and the writing process. N.D. conceptualized the study. All authors read, reviewed and approved the manuscript.

## Declaration of generative AI and AI-assisted technologies in the manuscript preparation process

During the preparation of this work, the author(s) used ChatGPT and Gemini on a limited basis to check grammar and readability of selected sentences. After using these tools, the author(s) reviewed and edited the content as needed and take(s) full responsibility for the content of the publication.

## References

1. MacDonald, M.E., Ambrose, C.M., Duyao, M.P., Myers, R.H., Lin, C., Srinidhi, L., Barnes, G., Taylor, S.A., James, M., Groot, N., et al. (1993). A novel gene containing a trinucleotide repeat that is expanded and unstable on Huntington’s disease chromosomes. The Huntington’s Disease Collaborative Research Group. Cell 72, 971–83.

2. Tabrizi, S.J., Schobel, S., Gantman, E.C., Mansbach, A., Borowsky, B., Konstantinova, P., Mestre, T.A., Panagoulias, J., Ross, C.A., Zauderer, M., et al. (2022). A biological classification of Huntington’s disease: the Integrated Staging System. Lancet Neurol 21, 632–644.

3. McColgan, P., and Tabrizi, S.J. (2018). Huntington’s disease: a clinical review. Eur J Neurol 25, 24–34.

4. Andhale, R., and Shrivastava, D. (2022). Huntington’s Disease: A Clinical Review. Cureus 14, e28484.

5. Benn, C.L., Sun, T., Sadri-Vakili, G., McFarland, K.N., DiRocco, D.P., Yohrling, G.J., Clark, T.W., Bouzou, B., and Cha, J.-H.J. (2008). Huntingtin modulates transcription, occupies gene promoters in vivo, and binds directly to DNA in a polyglutamine-dependent manner. J Neurosci 28, 10720–33.

6. Quintanilla, R.A., Jin, Y.N., Bernhardi, R.v., and Johnson, G.V.W. (2013). Mitochondrial permeability transition pore induces mitochondria injury in Huntington disease. Mol Neurodegener 8, 45.

7. Yano, H., Baranov, S.V., Baranova, O.V., Kim, J., Pan, Y., Yablonska, S., Carlisle, D.L., Ferrante, R.J., Kim, A.H., and Friedlander, R.M. (2014). Inhibition of mitochondrial protein import by mutant huntingtin. Nat Neurosci 17, 822–31.

8. Adegbuyiro, A., Sedighi, F., Pilkington, A.W., Groover, S., and Legleiter, J. (2017). Proteins Containing Expanded Polyglutamine Tracts and Neurodegenerative Disease. Biochemistry 56, 1199–1217.

9. Waldvogel, H.J., Kim, E.H., Tippett, L.J., Vonsattel, J.-P.G., and Faull, R.L.M. (2015). The Neuropathology of Huntington’s Disease. Curr Top Behav Neurosci 22, 33–80.

10. Goldberg, Y.P., Nicholson, D.W., Rasper, D.M., Kalchman, M.A., Koide, H.B., Graham, R.K., Bromm, M., Kazemi-Esfarjani, P., Thornberry, N.A., Vaillancourt, J.P., et al. (1996). Cleavage of huntingtin by apopain, a proapoptotic cysteine protease, is modulated by the polyglutamine tract. Nat Genet 13, 442–9.

11. Lunkes, A., Lindenberg, K.S., Ben-Haïem, L., Weber, C., Devys, D., Landwehrmeyer, G.B., Mandel, J.-L., and Trottier, Y. (2002). Proteases acting on mutant huntingtin generate cleaved products that differentially build up cytoplasmic and nuclear inclusions. Mol Cell 10, 259–69.

12. Gafni, J., Hermel, E., Young, J.E., Wellington, C.L., Hayden, M.R., and Ellerby, L.M. (2004). Inhibition of calpain cleavage of huntingtin reduces toxicity: accumulation of calpain/caspase fragments in the nucleus. J Biol Chem 279, 20211–20.

13. Graham, R.K., Deng, Y., Slow, E.J., Haigh, B., Bissada, N., Lu, G., Pearson, J., Shehadeh, J., Bertram, L., Murphy, Z., et al. (2006). Cleavage at the caspase-6 site is required for neuronal dysfunction and degeneration due to mutant huntingtin. Cell 125, 1179–91.

14. Chan, L.L., and Leavitt, B.R. (2026). Wild-type huntingtin in neurodevelopment. J Huntingtons Dis 15, 252–262.

15. (2025). Genetic modifiers of somatic expansion and clinical phenotypes in Huntington’s disease highlight shared and tissue-specific effects. Nat Genet 57, 1426– 1436.

16. Donaldson, J., Moss, D.H., Ciosi, M., Usdin, K., Balmus, G., Monckton, D.G., and Tabrizi, S.J. (2026). Huntington disease: somatic expansion, pathobiology and therapeutics. Nat Rev Neurol 22, 5–21.

17. E., H.R., Seva, K., Nora, R., Steven, T., Won-Seok, L., M., M.T., Kiely, M., Nolan, K., D., M.C., R., M.N., et al. (2025). Long somatic DNA-repeat expansion drives neurodegeneration in Huntington’s disease. Cell 188, 623–639.e19.

18. Kashin, S., Lee, W.-S., McDonald, T.M., Morris, K., Handsaker, R.E., Mello, C., Spina, L., Reed, N.M., Rivera, H.d., Jana, S.H., et al. (2025). A shared DNA-repeat toxicity threshold, reached somatically at cell-type-specific rates, unites cortical and striatal neurodegeneration in Huntington’s disease. Preprint at bioRxiv.

19. Pinto, R.M., Arning, L., Giordano, J.V., Razghandi, P., Andrew, M.A., Gillis, T., Correia, K., Mysore, J.S., Urtubey, D.-M.G., Parwez, C.R., et al. (2020). Patterns of CAG repeat instability in the central nervous system and periphery in Huntington’s disease and in spinocerebellar ataxia type 1. Hum Mol Genet 29, 2551–2567.

20. Barbaro, B.A., Lukacsovich, T., Agrawal, N., Burke, J., Bornemann, D.J., Purcell, J.M., Worthge, S.A., Caricasole, A., Weiss, A., Song, W., et al. (2015). Comparative study of naturally occurring huntingtin fragments in Drosophila points to exon 1 as the most pathogenic species in Huntington’s disease. Hum Mol Genet 24, 913–25.

21. Bragg, R.M., Landles, C., Smith, E.J., Osborne, G.F., Mathews, E.W., Cantle, J.P., Bates, G.P., and Carroll, J.B. (2026). Selective targeting of mutant huntingtin intron 1 improves rescue provided by antisense oligonucleotides in Huntington’s disease mice. Sci Transl Med 18, eadv0702.

22. Papadopoulou, A.S., Landles, C., Smith, E.J., Bondulich, M.K., Boeddrich, A., Canibano-Pico, M., Danby, E.C.E., Hoschek, F., Iqbal, A., Jones, S.T., et al. (2026). The HTT1a protein initiates HTT aggregation in a knock-in mouse model of Huntington’s disease. Brain.

23. Sathasivam, K., Neueder, A., Gipson, T.A., Landles, C., Benjamin, A.C., Bondulich, M.K., Smith, D.L., Faull, R.L.M., Roos, R.A.C., Howland, D., et al. (2013). Aberrant splicing of HTT generates the pathogenic exon 1 protein in Huntington disease. Proc Natl Acad Sci U S A 110, 2366–70.

24. Neueder, A., Landles, C., Ghosh, R., Howland, D., Myers, R.H., Faull, R.L.M., Tabrizi, S.J., and Bates, G.P. (2017). The pathogenic exon 1 HTT protein is produced by incomplete splicing in Huntington’s disease patients. Sci Rep 7, 1307.

25. Neueder, A., Dumas, A.A., Benjamin, A.C., and Bates, G.P. (2018). Regulatory mechanisms of incomplete huntingtin mRNA splicing. Nat Commun 9, 3955.

26. Hoschek, F., Natan, J., Wagner, M., Sathasivam, K., Abdelmoez, A., Einem, B.v., Bates, G.P., Landwehrmeyer, G.B., and Neueder, A. (2024). Huntingtin HTT1a is generated in a CAG repeat-length-dependent manner in human tissues. Mol Med 30, 36.

27. Mason, M.A., Gomez-Paredes, C., Sathasivam, K., Neueder, A., Papadopoulou, A.-S., and Bates, G.P. (2020). Silencing Srsf6 does not modulate incomplete splicing of the huntingtin gene in Huntington’s disease models. Sci Rep 10, 14057.

28. Pupak, A., Rodríguez-Navarro, I., Sathasivam, K., Singh, A., Essmann, A., Toro, D.D., Ginés, S., Pinto, R.M., Bates, G.P., Ørom, U.A.V., et al. (2024). m6A modification of mutant huntingtin RNA promotes the biogenesis of pathogenic huntingtin transcripts. EMBO Rep 25, 5026–5052.

29. Bai, D., Deng, F., Jia, Q., Ou, K., Wang, X., Hou, J., Zhu, L., Guo, M., Yang, S., Jiang, G., et al. (2024). Pathogenic TDP-43 accelerates the generation of toxic exon1 HTT in Huntington’s disease knock-in mice. Aging Cell 23, e14325.

30. Maffezzini, C., Iennaco, R., Scolz, A., Maestri, S., Landles, C., Osborne, G.F., Bronzini, M.M., Trovesi, C., Carzaniga, T., Casiraghi, L., et al. (2026). Human-specific sequence features in HTT exon 1 promote toxic misprocessing via splicing factor SRSF7. Nat Commun 17.

31. Mezer, M.d., Wojciechowska, M., Napierala, M., Sobczak, K., and Krzyzosiak, W.J. (2011). Mutant CAG repeats of Huntingtin transcript fold into hairpins, form nuclear foci and are targets for RNA interference. Nucleic Acids Res 39, 3852–63.

32. Green, K.M., Linsalata, A.E., and Todd, P.K. (2016). RAN translation-What makes it run? Brain Res 1647, 30–42.

33. Nguyen, L., Cleary, J.D., and Ranum, L.P.W. (2019). Repeat-Associated Non-ATG Translation: Molecular Mechanisms and Contribution to Neurological Disease. Annu Rev Neurosci 42, 227–247.

34. Bañez-Coronel, M., Ayhan, F., Tarabochia, A.D., Zu, T., Perez, B.A., Tusi, S.K., Pletnikova, O., Borchelt, D.R., Ross, C.A., Margolis, R.L., et al. (2015). RAN Translation in Huntington Disease. Neuron 88, 667–77.

35. Banez-Coronel, M., Zu, T., Aldridge, M., Guo, S., Ajredini, R., Morrison, D., Tays, A.B., Duvick, L.A., Pletnikova, O., Yachnis, A.T., et al. (2026). Repeat-associated non-AUG translation as a common mechanism for the polyGln ataxias. Cell Rep 45, 116741.

36. Tabrizi, S.J., Leavitt, B.R., Landwehrmeyer, G.B., Wild, E.J., Saft, C., Barker, R.A., Blair, N.F., Craufurd, D., Priller, J., Rickards, H., et al. (2019). Targeting Huntingtin Expression in Patients with Huntington’s Disease. N Engl J Med 380, 2307–2316.

37. McColgan, P., Thobhani, A., Boak, L., Schobel, S.A., Nicotra, A., Palermo, G., Trundell, D., Zhou, J., Schlegel, V., Ducray, P.S., et al. (2023). Tominersen in Adults with Manifest Huntington’s Disease. N Engl J Med 389, 2203–2205.

38. Bas, R., Janwillem, T., Rudie, W., Chantal, B., and A., D.N. (2025). Accurate Quantification of Mutant and Wild-Type polyQ Proteins Using Simple Western Capillary Immunoassays. Mol Neurobiol 62, 12768–12790.

39. Mathews, E.W., Coffey, S.R., Gärtner, A., Belgrad, J., Bragg, R.M., O’Reilly, D., Cantle, J.P., McHugh, C., Summers, A., Fentz, J., et al. (2025). Suppression of Huntington’s Disease Somatic Instability by Transcriptional Repression and Direct CAG Repeat Binding. Nat Commun 16, 10009.

40. Wheeler, V.C., Auerbach, W., White, J.K., Srinidhi, J., Auerbach, A., Ryan, A., Duyao, M.P., Vrbanac, V., Weaver, M., Gusella, J.F., et al. (1999). Length-dependent gametic CAG repeat instability in the Huntington’s disease knock-in mouse. Hum Mol Genet 8, 115–22.

41. Missineo, A., Tomei, L., Alaimo, N., Martufi, P., Zavattieri, M., Colicchia, V., Cariulo, C., Fodale, V., Séguin, J., Esquina, C., et al. (2026). Development and validation of highly selective monoclonal antibodies for the detection of huntingtin neoepitopes. Preprint at bioRxiv.

42. Davies, S.W., Turmaine, M., Cozens, B.A., DiFiglia, M., Sharp, A.H., Ross, C.A., Scherzinger, E., Wanker, E.E., Mangiarini, L., and Bates, G.P. (1997). Formation of neuronal intranuclear inclusions underlies the neurological dysfunction in mice transgenic for the HD mutation. Cell 90, 537–48.

43. Papadopoulou, A.S., Alterman, J., Landles, C., Smith, E.J., Conroy, F., Phillips, J., Canibano-Pico, M., Nita, I.M., Osborne, G.F., Iqbal, A., et al. (2026). Lowering the HTT1a transcript as an effective therapy for Huntington’s disease in a knockin mouse model. Sci Transl Med 18, eadw2495.

44. Castelli, L.M., Huang, W.-P., Lin, Y.-H., Chang, K.-Y., and Hautbergue, G.M. (2021). Mechanisms of repeat-associated non-AUG translation in neurological microsatellite expansion disorders. Biochem Soc Trans 49, 775–792.

45. Krans, A., Albertson, R.M., Maltby, C.J., and Todd, P.K. (2026). Mechanistic insights into RAN translation within the Huntingtin coding sequence. Preprint at bioRxiv.

46. Jazurek-Ciesiolka, M., Ciesiolka, A., Komur, A.A., Urbanek-Trzeciak, M.O., Krzyzosiak, W.J., and Fiszer, A. (2020). RAN Translation of the Expanded CAG Repeats in the SCA3 Disease Context. J Mol Biol 432, 166699.

47. Crooke, S.T., Liang, X.-H., Baker, B.F., and Crooke, R.M. (2021). Antisense technology: A review. J Biol Chem 296, 100416.

48. Cattaneo, E. (2003). Dysfunction of wild-type huntingtin in Huntington disease. News Physiol Sci 18, 34–7.

49. Schulte, J., and Littleton, J.T. (2011). The biological function of the Huntingtin protein and its relevance to Huntington’s Disease pathology. Curr Trends Neurol 5, 65–78.

50. Dietrich, P., Johnson, I.M., Alli, S., and Dragatsis, I. (2017). Elimination of huntingtin in the adult mouse leads to progressive behavioral deficits, bilateral thalamic calcification, and altered brain iron homeostasis. PLoS Genet 13, e1006846.

51. Bragg, R.M., Mathews, E.W., Grindeland, A., Cantle, J.P., Howland, D., Vogt, T., and Carroll, J.B. (2024). Global huntingtin knockout in adult mice leads to fatal neurodegeneration that spares the pancreas. Life Sci Alliance 7.

52. Raamsdonk, J.M.V., Pearson, J., Rogers, D.A., Bissada, N., Vogl, A.W., Hayden, M.R., and Leavitt, B.R. (2005). Loss of wild-type huntingtin influences motor dysfunction and survival in the YAC128 mouse model of Huntington disease. Hum Mol Genet 14, 1379–92.

53. McBride, J.L., Pitzer, M.R., Boudreau, R.L., Dufour, B., Hobbs, T., Ojeda, S.R., and Davidson, B.L. (2011). Preclinical safety of RNAi-mediated HTT suppression in the rhesus macaque as a potential therapy for Huntington’s disease. Mol Ther 19, 2152– 62.

54. Ambrose, C.M., Duyao, M.P., Barnes, G., Bates, G.P., Lin, C.S., Srinidhi, J., Baxendale, S., Hummerich, H., Lehrach, H., Altherr, M., et al. (1994). Structure and expression of the Huntington’s disease gene: evidence against simple inactivation due to an expanded CAG repeat. Somat Cell Mol Genet 20, 27–38.

55. Ciosi, M., Maxwell, A., Cumming, S.A., Moss, D.J.H., Alshammari, A.M., Flower, M.D., Durr, A., Leavitt, B.R., Roos, R.A.C., et al. (2019). A genetic association study of glutamine-encoding DNA sequence structures, somatic CAG expansion, and DNA repair gene variants, with Huntington disease clinical outcomes. EBioMedicine 48, 568–580.

56. Roy, J.C.L., Vitalo, A., Andrew, M.A., Mota-Silva, E., Kovalenko, M., Burch, Z., Nhu, A.M., Cohen, P.E., Grabczyk, E., Wheeler, V.C., et al. (2021). Somatic CAG expansion in Huntington’s disease is dependent on the MLH3 endonuclease domain, which can be excluded via splice redirection. Nucleic Acids Res 49, 3907–3918.

57. Lee, J.-M., Huang, Y., Orth, M., Gillis, T., Siciliano, J., Hong, E., Mysore, J.S., Lucente, D., Wheeler, V.C., Seong, I.S., et al. (2022). Genetic modifiers of Huntington disease differentially influence motor and cognitive domains. Am J Hum Genet 109, 885–899.

58. Wheeler, V.C., and Dion, V. (2021). Modifiers of CAG/CTG Repeat Instability: Insights from Mammalian Models. J Huntingtons Dis 10, 123–148.

59. Sadler, F., and Tabrizi, S.J. (2026). Targeting toxic HTT1a could be the key to treating Huntington disease. Nat Rev Neurol 22, 547–548.

60. Scahill, R.I., Farag, M., Murphy, M.J., Hobbs, N.Z., Leocadi, M., Langley, C., Knights, H., Ciosi, M., Fayer, K., Nakajima, M., et al. (2025). Somatic CAG repeat expansion in blood associates with biomarkers of neurodegeneration in Huntington’s disease decades before clinical motor diagnosis. Nat Med 31, 807–818.

61. Datson, N.A., González-Barriga, A., Kourkouta, E., Weij, R., Giessen, J.v.d., Mulders, S., Kontkanen, O., Heikkinen, T., Lehtimäki, K., and Deutekom, J.C.T.v. (2017). The expanded CAG repeat in the huntingtin gene as target for therapeutic RNA modulation throughout the HD mouse brain. PLoS One 12, e0171127.

62. Girstmair, H., Saffert, P., Rode, S., Czech, A., Holland, G., Bannert, N., and Ignatova, Z. (2013). Depletion of cognate charged transfer RNA causes translational frameshifting within the expanded CAG stretch in huntingtin. Cell Rep 3, 148–59.

63. Wojciechowska, M., Olejniczak, M., Galka-Marciniak, P., Jazurek, M., and Krzyzosiak, W.J. (2014). RAN translation and frameshifting as translational challenges at simple repeats of human neurodegenerative disorders. Nucleic Acids Res 42, 11849–64.

64. Saffert, P., Adamla, F., Schieweck, R., Atkins, J.F., and Ignatova, Z. (2016). An Expanded CAG Repeat in Huntingtin Causes +1 Frameshifting. J Biol Chem 291, 18505–13.

65. Chung, D.W., Rudnicki, D.D., Yu, L., and Margolis, R.L. (2011). A natural antisense transcript at the Huntington’s disease repeat locus regulates HTT expression. Hum Mol Genet 20, 3467–77.

66. Rué, L., Bañez-Coronel, M., Creus-Muncunill, J., Giralt, A., Alcalá-Vida, R., Mentxaka, G., Kagerbauer, B., Zomeño-Abellán, M.T., Aranda, Z., Venturi, V., et al. (2016). Targeting CAG repeat RNAs reduces Huntington’s disease phenotype independently of huntingtin levels. J Clin Invest 126, 4319–4330.

67. Nalavade, R., Griesche, N., Ryan, D.P., Hildebrand, S., and Krauss, S. (2013). Mechanisms of RNA-induced toxicity in CAG repeat disorders. Cell Death Dis 4, e752.

68. Martí, E. (2016). RNA toxicity induced by expanded CAG repeats in Huntington’s disease. Brain Pathol 26, 779–786.

69. Heinz, A., Nabariya, D.K., and Krauss, S. (2021). Huntingtin and Its Role in Mechanisms of RNA-Mediated Toxicity. Toxins 13.

70. Thomas, J.D., Oliveira, R., Sznajder, Ł.J., and Swanson, M.S. (2018). Myotonic Dystrophy and Developmental Regulation of RNA Processing. Compr Physiol 8, 509– 553.

71. McEachin, Z.T., Parameswaran, J., Raj, N., Bassell, G.J., and Jiang, J. (2020). RNA-mediated toxicity in C9orf72 ALS and FTD. Neurobiol Dis 145, 105055.

72. Dincã, D.M., Lallemant, L., González-Barriga, A., Cresto, N., Braz, S.O., Sicot, G., Pillet, L.-E., Polvèche, H., Magneron, P., Huguet-Lachon, A., et al. (2022). Myotonic dystrophy RNA toxicity alters morphology, adhesion and migration of mouse and human astrocytes. Nat Commun 13, 3841.

73. Goodwin, M., Mohan, A., Batra, R., Lee, K.-Y., Charizanis, K., Gómez, F.J.F., Eddarkaoui, S., Sergeant, N., Buée, L., Kimura, T., et al. (2015). MBNL Sequestration by Toxic RNAs and RNA Misprocessing in the Myotonic Dystrophy Brain. Cell Rep 12, 1159–68.

74. Lee, Y.-B., Chen, H.-J., Peres, J.N., Gomez-Deza, J., Attig, J., Stalekar, M., Troakes, C., Nishimura, A.L., Scotter, E.L., Vance, C., et al. (2013). Hexanucleotide repeats in ALS/FTD form length-dependent RNA foci, sequester RNA binding proteins, and are neurotoxic. Cell Rep 5, 1178–86.

75. Ly, S., Didiot, M.-C., Ferguson, C.M., Coles, A.H., Miller, R., Chase, K., Echeverria, D., Wang, F., Sadri-Vakili, G., Aronin, N., et al. (2022). Mutant huntingtin messenger RNA forms neuronal nuclear clusters in rodent and human brains. Brain Commun 4, fcac248.

76. Jafar-Nejad, P., Powers, B., Soriano, A., Zhao, H., Norris, D.A., Matson, J., DeBrosse-Serra, B., Watson, J., Narayanan, P., Chun, S.J., et al. (2021). The atlas of RNase H antisense oligonucleotide distribution and activity in the CNS of rodents and non-human primates following central administration. Nucleic Acids Res 49, 657–673.

77. Luu, K.T., Norris, D.A., Gunawan, R., Henry, S., Geary, R., and Wang, Y. (2017). Population Pharmacokinetics of Nusinersen in the Cerebral Spinal Fluid and Plasma of Pediatric Patients With Spinal Muscular Atrophy Following Intrathecal Administrations. J Clin Pharmacol 57, 1031–1041.

78. Rosas, H.D., Salat, D.H., Lee, S.Y., Zaleta, A.K., Pappu, V., Fischl, B., Greve, D., Hevelone, N., and Hersch, S.M. (2008). Cerebral cortex and the clinical expression of Huntington’s disease: complexity and heterogeneity. Brain 131, 1057–68.

79. Unschuld, P.G., Joel, S.E., Liu, X., Shanahan, M., Margolis, R.L., Biglan, K.M., Bassett, S.S., Schretlen, D.J., Redgrave, G.W., Zijl, P.C.M.v., et al. (2012). Impaired cortico-striatal functional connectivity in prodromal Huntington’s Disease. Neurosci Lett 514, 204–9.

80. Bunner, K.D., and Rebec, G.V. (2016). Corticostriatal Dysfunction in Huntington’s Disease: The Basics. Front Hum Neurosci 10, 317.

81. Blumenstock, S., and Dudanova, I. (2020). Cortical and Striatal Circuits in Huntington’s Disease. Front Neurosci 14, 82.

82. Johnson, E.B., Ziegler, G., Penny, W., Rees, G., Tabrizi, S.J., Scahill, R.I., and Gregory, S. (2021). Dynamics of Cortical Degeneration Over a Decade in Huntington’s Disease. Biol Psychiatry 89, 807–816.

83. Paepe, A.E.D., Bikou, V., Turan, E., Pérez-Bellido, A., Garcia-Gorro, C., Rodriguez-Dechicha, N., Vaquer, I., Calopa, M., Diego-Balaguer, R.d., and Camara, E. (2025). Striato-cortical connectivity patterns predict clinical profiles in Huntington’s disease. Neuroimage Clin 46, 103788.

84. Hammond, S.M., Abendroth, F., Goli, L., Stoodley, J., Burrell, M., Thom, G., Gurrell, I., Ahlskog, N., Gait, M.J., Wood, M.J., et al. (2022). Antibody-oligonucleotide conjugate achieves CNS delivery in animal models for spinal muscular atrophy. JCI Insight 7.

85. Barker, S.J., Thayer, M.B., Kim, C., Tatarakis, D., Simon, M.J., Dial, R., Nilewski, L., Wells, R.C., Zhou, Y., Afetian, M., et al. (2024). Targeting the transferrin receptor to transport antisense oligonucleotides across the mammalian blood-brain barrier. Sci Transl Med 16, eadi2245.

86. Iwamoto, N., Liu, Y., Frank-Kamenetsky, M., Maguire, A., Tseng, W.C., Taborn, K., Kothari, N., Akhtar, A., Bowman, K., Shelke, J.D., et al. (2024). Preclinical evaluation of stereopure antisense oligonucleotides for allele-selective lowering of mutant HTT. Mol Ther Nucleic Acids 35, 102246.

87. Gonzalez, E.V., and Zoghbi, H.Y. (2026). Pathogenesis of polyglutamine diseases: Piecing together a complex molecular puzzle. J Exp Med 223.

88. Kourkouta, E., Weij, R., González-Barriga, A., Mulder, M., Verheul, R., Bosgra, S., Groenendaal, B., Puoliväli, J., Toivanen, J., Deutekom, J.C.T.v., et al. (2019). Suppression of Mutant Protein Expression in SCA3 and SCA1 Mice Using a CAG Repeat-Targeting Antisense Oligonucleotide. Mol Ther Nucleic Acids 17, 601–614.

89. Farag, M., Tabrizi, S.J., and Wild, E.J. (2026). Huntington’s disease clinical trials update: October 2025. J Huntingtons Dis 15, 156–166.

90. DeVos, S.L., and Miller, T.M. (2013). Direct intraventricular delivery of drugs to the rodent central nervous system. J Vis Exp, e50326.

91. Lee, J.-M., Zhang, J., Su, A.I., Walker, J.R., Wiltshire, T., Kang, K., Dragileva, E., Gillis, T., Lopez, E.T., Boily, M.-J., et al. (2010). A novel approach to investigate tissue-specific trinucleotide repeat instability. BMC Syst Biol 4, 29.

