## Supplemental_Figures S1-S4_and_Tables S1-4 for "VO659, a CAG repeat-targeting antisense oligonucleotide, lowers multiple neurotoxic mutant huntingtin proteins in Huntington’s disease"

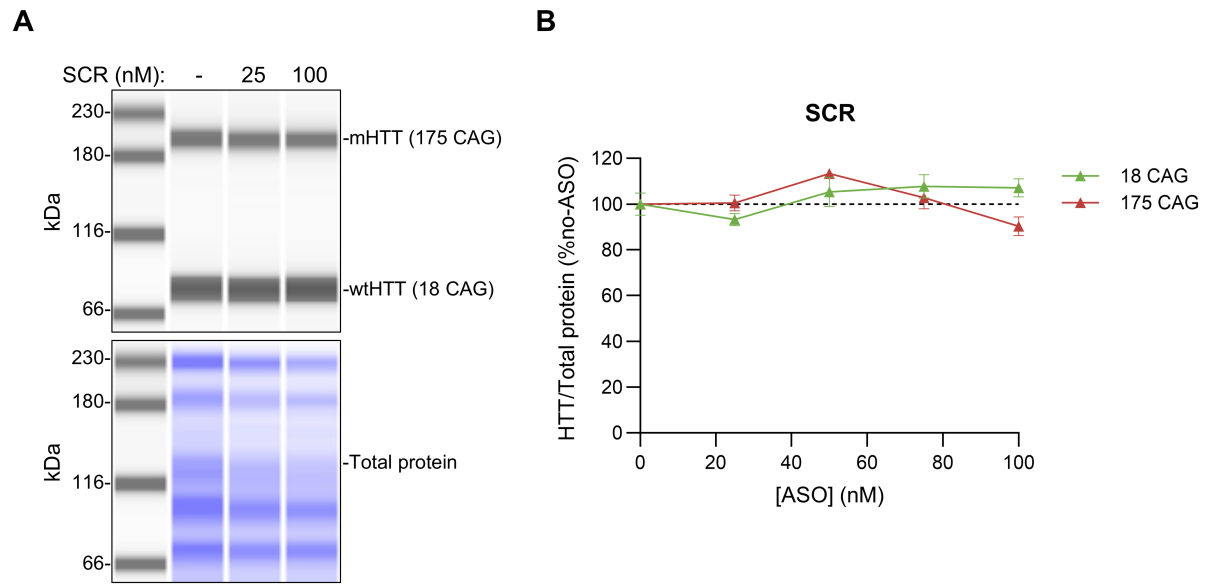

**Supplemental Figure S1** A scrambled version of VO659 (SCR) does not decrease endogenous HTT levels **A** Representative Simple Western (SW) blot view of HD patient fibroblasts with 175:18 CAGs in *HTT* treated with 25–100 nM of a scrambled version of VO659 (SCR). To enable separate separation of wtHTT and mHTT, protein lysate was treated with 6 μg/mL caspase-3 (Bio-Techne, Minneapolis, MN, 707-C3/CF) for 1 hour at 37 °C.<sup>38</sup> Then, 0.21 mg/mL of the treated protein lysate was loaded on 12–230 kDa SW module plates. HTT (Abcam, ab109115, 1:100) and total protein (for normalization) were detected as described in the materials and methods section. **B** Quantification of HTT protein levels in HD patient fibroblasts transfected with SCR. No effect of SCR was observed on wtHTT and mHTT protein levels. HTT levels are expressed as % of no-ASO control after correction for total protein. The mean ± SD of three culture wells from a single transfection experiment is shown.

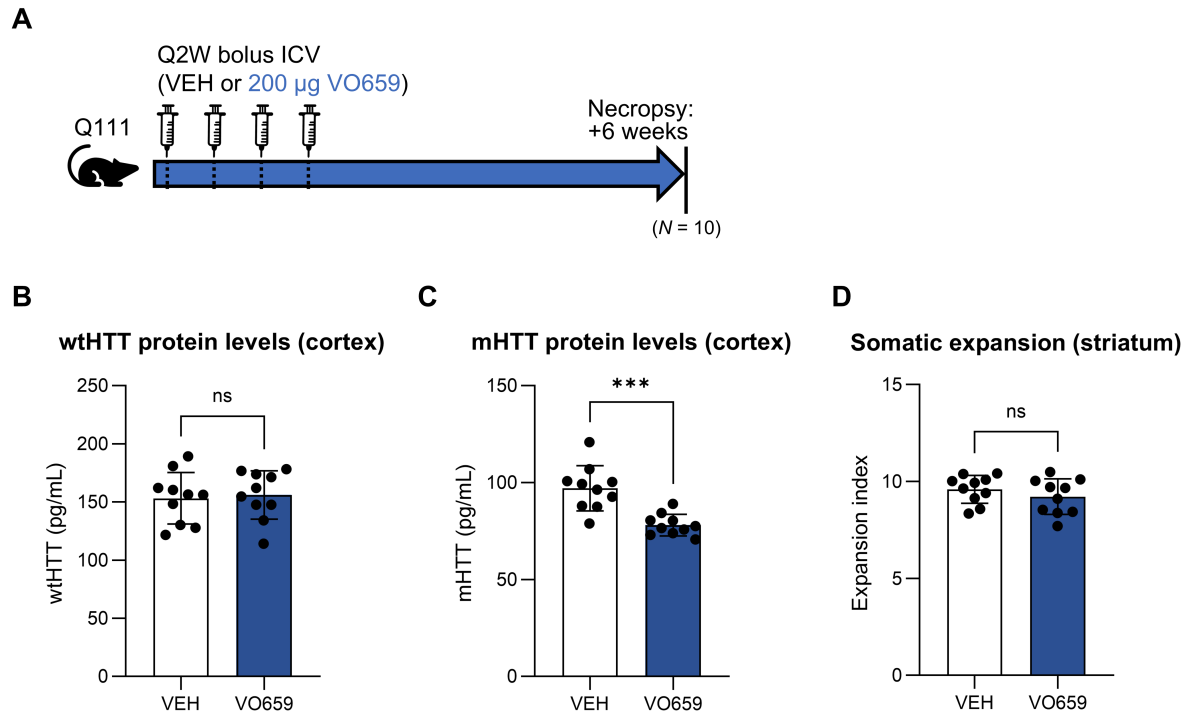

**Supplemental Figure S2** VO659 does not affect somatic expansion in striatum of Q111 mice **A** Outline of an *in vivo* study in Q111 HD mice. Mice were given a total of 4 ICV bolus doses of 200 µg of VO659 or vehicle (VEH) once per 2 weeks (Q2W), starting at the age of 2.5 months ($N = 10$). The mice were sacrificed 6 weeks after the last dose. Cortex was collected for quantification of wtHTT and mHTT protein levels, while striatum was collected to determine the somatic expansion index. **B–C** wtHTT and mHTT protein levels as quantified with Meso Scale Discovery immunoassays (MSD) in cortex tissue **D** Somatic expansion index of the CAG repeat in Q111 mice striatum. Data are expressed as the mean ± SD. Groups were compared using unpaired T-tests using GraphPad Prism. \* = $p < 0.05$, \*\* = $p < 0.005$, \*\*\* = $p < 0.0005$, ns = not significant

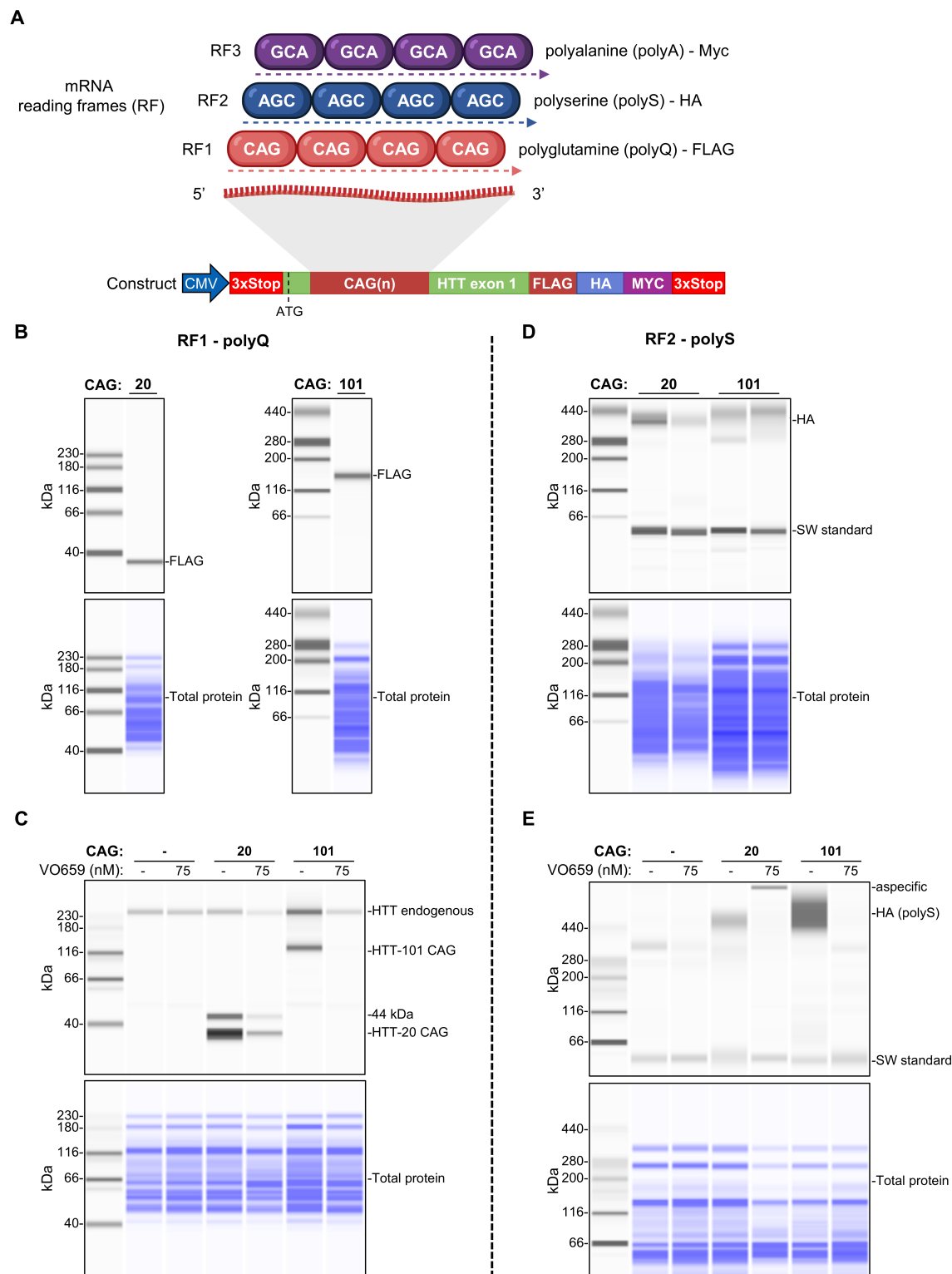

**Supplemental Figure S3** *PolyQ and polyS RAN translation was not consistently detectable* **A** Schematic of out-of-frame translation from a *HTT* expression construct with 20 or 101 CAG repeats containing tags in the three different reading frames (same as in Fig. 5A). Translation from reading frame 1 (RF1), including canonical translation, results in expression of FLAG-tagged polyglutamine (polyQ) containing proteins, translation from RF2 results in HA-tagged polyserine (polyS) containing proteins, and translation from RF3 results in Myc-tagged polyaniline (polyA) containing proteins. **B–E** Representative Simple Western (SW) blot views of HeLa cells transfected with the construct described in A and, when indicated, 75 nM VO659 (**B**) SW runs probed with an anti-FLAG tag antibody (Bio-Techne, #MAB8529, Minneapolis, 1:100). No high molecular weight (Mw) polyQ smear was detected.

Only 2 distinct bands for the 20 and 101 CAG repeat constructs were observed, corresponding to the canonical open reading frame (ORF) protein product (**C**) SW run probed with an antibody targeting the N-terminus of HTT (Abcam, #ab109115, 1:100). Bands corresponding to endogenous HTT from the HeLa cells, and bands corresponding to the canonical ORF proteins from the 20 and 101 CAG repeat constructs were observed. The 20 and 101 CAG repeat construct protein bands were very similar to the bands observed in B, suggesting that protein product translated from RF1 from the constructs contained both the HTT N-terminus and the FLAG tag, and are therefore not RAN translation-derived. A 44 kDa band of unknown origin was detected in cells transfected with the 20 CAG construct. It is unlikely this is a RAN translation product as a similar band was not detected with 101 CAG repeats (**D**) SW run probed with an anti-HA tag antibody (Bio-Techne, #MAB0601, 1:25). HA signal was very low, suggesting either low abundance or poor detection of a potentially aggregated polyS RAN translation-derived protein. The low Mw band corresponds to a SW fluorescent standard added to every sample during an SW run, which is shown very prominently here due to other signal being low (**E**) SW run probed with an anti-HA tag antibody (Bio-Techne, #MAB0601, 1:25). In this experiment, HA-tagged polyS signal from RF2 could be detected as a high Mw smear which increased in intensity and Mw with CAG repeat length, although still at low levels. Transfection with VO659 reduced HA signal. For all blots, a Total protein stain is shown as control. Note, for figures **B** and **D**, the transfection protocol as outlined in the Materials and methods section and Supplemental Figure S4 was used. For **C** and **E**, the order of transfection as outlined in Supplemental Figure S4 was swapped, meaning ASO was transfected first, followed by plasmid.

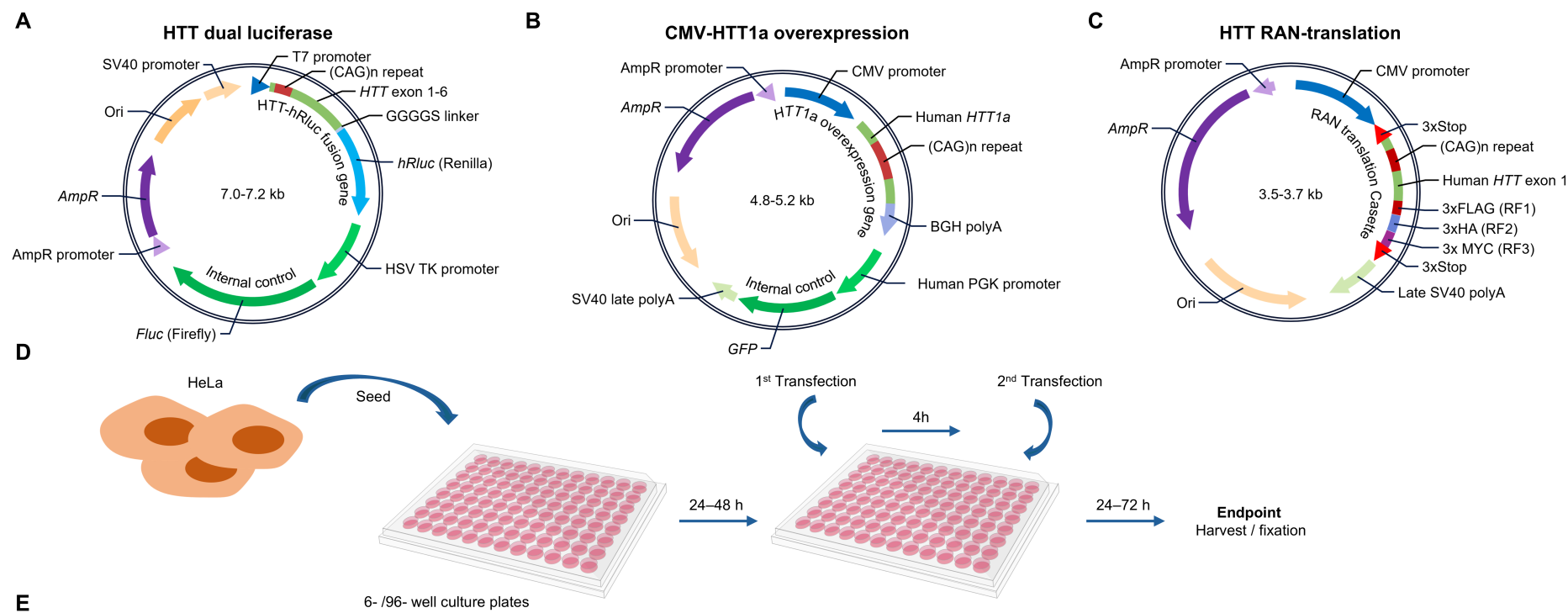

| Cell line | Construct | Plate type | Seed density | Seeded (h before transfection) | Endpoint (h after transfection) | Transfection volume |  |  | 1<sup>st</sup> Transfection |  | 2<sup>nd</sup> Transfection |  |
| --- | --- | --- | --- | --- | --- | --- | --- | --- | --- | --- | --- | --- |
|  |  |  |  |  |  | MEM (μL) | Opti-MEM (μL) | Total (μL) | Payload | Reagent | Payload | Reagent |
| HeLa | HTT dual luciferase | White 96 well plate (Greiner Bio-One, #655083) | 12500 | 24 | 24 | 37.5 | 37.5 | 75 | 0.5–60 nM ASO | 0.1 μL Lipofectamine-2000 | 80 ng Plasmid | 0.2 μL Lipofectamine-2000 |
| HeLa | HTT1a overexpression | Black 96 well plate (Corning, #CLS3603) | 10000 | 24 | 72 | 37.5 | 37.5 | 75 | 5–50 nM ASO | 0.1 μL Lipofectamine RNAiMAX | 60 ng Plasmid | 0.1 μL Lipofectamine-3000 |
| HeLa | HTT RAN-translation | 6 well plate (Corning, #734-1599) | 200000 - 250000 | 48 | 24 | 1000 | 250 | 1250 | 1000 ng Plasmid | 5 μL Lipofectamine-2000 | 35–75 nM ASO | 0.7–1.5 μL Lipofectamine RNAiMAX |

**Supplemental Figure S4. Co-transfection protocols of ASO and plasmid in HeLa cells. A–C** Schematics of plasmids used in double transfection experiments. **A** HTT dual luciferase reporter (DLR) construct containing *HTT* exon 1–6 with 5–98 CAG repeats fused to a Renilla luciferase (*hRluc*) using a GGGGS flexible linker. An internal control Firefly luciferase (*fLuc*) is driven under a separate HSV TK promoter. The constructs were generated in house using the psiCHEK-2 backbone (Promega, Madison, WI, #C8021) **B** The plasmid used in *HTT1a* overexpression experiments containing a *HTT1a* (100 or 156 CAG repeats)

expression cassette under a control of a CMV promoter and a *GFP* internal reference gene under control of a human PGK promoter, generated in-house using a custom pRP backbone obtained from VectorBuilder (Chicago, IL) **C** Plasmid used for RAN translation experiments containing a *HTT* exon 1 expression construct with 20 or 100 CAG repeats under a CMV promoter with triple FLAG, HA or Myc tags in the three different sense reading frames. The constructs also contained triple Stop cassettes 5' and 3' of the HTT construct. RAN translation plasmids were synthesized by VectorBuilder (Chicago, IL). All plasmids contained Origin of Replication (Ori, in yellow) and ampicillin resistance (*AmpR*, in purple) sequences. **D** Schematic workflow of double transfection experiments using HeLa cells. **E** Table showing transfection conditions per plasmid, including plate type, seeding density, timing of transfection and harvest, transfection volumes, order of transfection, transfection reagents, ASO concentrations, and plasmid amounts

### Supplemental tables

**Supplemental Table S1. List of ASOs**

| ASO | Chemistry | Sequence 5'→3' |
| --- | --- | --- |
| VO659 | 2'-O-Me PS | C*•U•G•C*•U•G•C*•U•G•C*•U•G•C*•U•G•C*•U•G |
| SCR | 2'-O-Me PS | C*•U•G•U•G•G•U•C*•C*•U•C*•C*•U•G•G•C*•C*•U•U•G•G |
| HTT-gap36 | 2'-MOE PO/PS gapmer | C*•T•C*•A•G•T•A•A•C*•A•T•T•G•A•C*•A•C*•C*•A•C* |

Blue: 2'-O-Methyl (2'-O-Me) sugar modification; Green: 2'-methoxyethyl (3'-MOE) sugar modification; Red: DNA, • = phosphorothioate (PS) backbone linkage, ○ = phosphodiester (PO) backbone linkage, \* = Methylated cytosine base. VO659 is also known as (CUG)7.<sup>61,88</sup> HTT-gap36 has the same sequence and chemistry as HTT<sub>Rx</sub>, also known as tominersen.<sup>36-37</sup>

**Supplemental Table S2. Simple Western (SW) antibodies and loading concentrations**

| Target | Antibody | Vendor | Dilution (v/v) | Host species | Clonality | Loading concentration (mg/mL protein) |
| --- | --- | --- | --- | --- | --- | --- |
| HTT n-terminus | #ab109115 | Abcam, Cambridge, UK | 1:100 | Rabbit | Monoclonal | 0.07* |
| FLAG tag | #MAB8529 | Bio-Techne, Minneapolis, MN | 1:100 | Rabbit | Monoclonal | 0.5** |
| HA tag | #MAB0601 | Bio-Techne, Minneapolis, MN | 1:25 | Rabbit | Monoclonal | 0.5** |
| Myc tag | #2278 | Cell Signaling Technology, Danvers, MA | 1:25 | Rabbit | Monoclonal | 0.5** |

\*0.21 mg/mL was loaded for Supplemental figure S1. \*\*0.5 mg/mL was the standard input for these SW experiments. However, due to high expression levels from plasmids, occasional SW signal saturation was observed in specific experiments. If saturated signal was observed, samples from that experiment were re-measured using a 0.1 mg/mL protein input.

**Supplemental Table S3. RT-dPCR primers and TaqMan probes**

| Target | Primer | Sequence 5'→3' |
| --- | --- | --- |
| Poly(A) | RT-oligo(dT)UAP* | GGCCACGCGTCGACTAGTACTTTTTTTTTTTTTTTTTT |
| HTT1a TaqMan assay | HTT1a_F Forward | GGATCCACACTCAAAACATTTA |
|  | HTT1a_probe (FAM)** | /6-FAM/TCTTATTC/ZEN/AGACAACAAGGAGGAAAAATAAAATACC/IABkFQ/ |
|  | UAP_R Reverse | CACGCGTCGACTAGTAC |
| HTT exon 2 TaqMan assay | HTT exon 2_F Forward | AAAGAAAGAACTTTCAGCTACCAAGAA |
|  | HTT exon 2_Probe (HEX)** | /HEX/TCACATATT/ZEN/GTCAGACAATGATTCACACGGTCT/IABkFQ/ |
|  | HTT exon 2_R Reverse | CTGACAGACTGTGCCACTATG |
| GAPDH | GAPDH assay (FAM) | Proprietary (#Hs04420632_g1, FAM-MGB) |

\*The primer used to generate cDNA in the reverse transcriptase (RT) reaction. \*\*For TaqMan probes, the fluorophore (6-FAM or HEX) is indicated in the sequence together with the double quenchers, ZEN and Iowa Black FQ (/3IABkFQ/). Custom primers and probes were synthesized by Integrated DNA technologies (IDT; Leuven, Belgium). The premade Hs04420632\_g1 GAPDH TaqMan assay was obtained from Thermo Fisher Scientific, Waltham, MA. The sequences for the RT-oligo(dT)UAP primer, HTT1a and HTT exon 2 and TaqMan assays were taken from Hoschek et al.<sup>26</sup>

**Supplemental Table S4. Sandwich Hybridization ELISA capture and detection probes**

| Probe | Target | Sequence 5'→3' |
| --- | --- | --- |
| Capture probe-1 | VO659 | Phosphate-CAGCAGCAG<u>**AG**</u>-biotin-TEG |
| Detection probe-1 | VO659 | Digoxigenin-<u>**CAGCAGCAG**</u> |
| Capture probe-2 | HTT-gap36 | Phosphate-ATGTTACTG<u>**AG**</u>-biotin-TEG |
| Detection probe-2 | HTT-gap36 | Digoxigenin-<u>**GTGGTGTCA**</u> |

ELISA probes were obtained from Eurogentec (Seraing, Belgium). Boldened underlined bases contained a locked nucleic acid (LNA) sugar-ring modification. The other bases were unmodified DNA.
